# Polygenic adaptation and negative selection across traits, years and environments in a long-lived plant species (*Pinus pinaster* Ait., Pinaceae)

**DOI:** 10.1101/2020.03.02.974113

**Authors:** Marina de Miguel, Isabel Rodríguez-Quilón, Myriam Heuertz, Agathe Hurel, Delphine Grivet, Juan-Pablo Jaramillo-Correa, Giovanni G. Vendramin, Christophe Plomion, Juan Majada, Ricardo Alía, Andrew J. Eckert, Santiago C. González-Martínez

## Abstract

A decade of association studies in multiple organisms suggests that most complex traits are polygenic; that is, they have a genetic architecture determined by numerous loci distributed across the genome, each with small effect-size. Thus, determining the degree of polygenicity and its variation across traits, environments and years is useful to understand the genetic basis of phenotypic variation. In this study, we applied multilocus approaches to estimate the degree of polygenicity of fitness-related traits in a long-lived plant (*Pinus pinaster* Ait., maritime pine) and to analyze how polygenicity changes across environments and years. To do so, we evaluated five categories of fitness-related traits (survival, height, phenology-related, functional, and biotic-stress response traits) in a clonal common garden network, planted in contrasted environments (over 12,500 trees). First, most of the analyzed traits showed evidence of local adaptation based on *Q*_ST_-*F*_ST_ comparisons. Second, we observed a remarkably stable degree of polygenicity, averaging 6% (range of 0-27%), across traits, environments and years. As previously suggested for humans, some of these traits showed also evidence of negative selection, which could explain, at least partially, the high degree of polygenicity. The observed genetic architecture of fitness-related traits in maritime pine supports the polygenic adaptation model. Because polygenic adaptation can occur rapidly, our study suggests that current predictions on the capacity of natural forest tree populations to adapt to new environments should be revised, which is of special relevance in the current context of climate change.

## Introduction

Population adaptive responses to environmental changes depend on the genetic architecture of fitness-related traits (Hayward and Sella 2019). Although not initially conceived for the study of adaptation, genome-wide association studies (GWAS) have provided essential information to understand the genetic basis of complex traits. The implementation of GWAS allowed the identification of genetic variants affecting fitness-related traits, their allele frequencies, the magnitudes of their effects, and their interactions with one another and the environment. Examples exist for humans (reviewed by Visscher *et al.* 2017), other animals (e.g. Sharma *et al.* 2015; Pitchers *et al.* 2019) and plants (e.g. González-Martínez *et al.* 2006; Huang and Han 2014; Alonso-Blanco *et al.* 2016). Surprisingly, in some species, such as humans and forest trees (Resende *et al.* 2012; Lind *et al.* 2018), the genetic variants associated with phenotypic variation accounted for only small fractions of trait heritability, as estimated through pedigree analysis, causing the so-called ‘missing heritability’ paradox (Maher 2008). Several explanations have been provided to solve this paradox (Manolio *et al.* 2009; Brachi *et al.* 2011; Björkegren *et al.* 2015; Pallares 2019). In particular, different sources of evidence point to polygenicity, i.e. trait architecture determined by a large number of variants, each with a small effect-size, as a potential reason for the low levels of heritability explained by current GWAS, which would thus be unpowered to detect most causal variants (Yang *et al.* 2010; Shi *et al.* 2016; Boyle *et al.* 2017).

The study of adaptation has traditionally been addressed from contrasting research paradigms (Höllinger *et al.* 2019). While quantitative genetic approaches view adaptation as the result of changes in allele frequencies at an idealized infinite number of loci, each with infinitesimal effects on fitness (Fisher 1918), population genetic approaches made more emphasis in the detection of selective sweeps, where new beneficial mutations rapidly become fixed at a small number of loci (Smith and Haigh 1974). The hypothesis that natural selection (mostly) acts through subtle allele frequency shifts on standing genetic variation at numerous loci distributed across the genome has been suggested in previous evolutionary studies (Orr and Coyne 1992; e.g. McKay and Latta 2002; Le Corre and Kremer 2003). Nevertheless, it was Pritchard *et al.* (2010) who first brought together population and quantitative genetic theory with conclusions from GWAS to formulate a new model for the study of adaptation – the polygenic adaptation model. Under this model, some genes may harbor new mutations that have been fixed by natural selection, but the most common pattern would be the genome-wide increase of favored alleles, without the fixation of most causative variants. Thus, the expected genome-wide footprint resulting from natural selection would not be that of a classical hard sweep, but would rather involve a large number of causal variants, each with subtle allele frequency changes (Pritchard *et al.* 2010; Pritchard and Rienzo 2010; Hermisson and Pennings 2017).

Several new methods have been developed to detect the genetic signatures of natural selection under the polygenic adaptation model (Guan and Stephens 2011; Turchin *et al.* 2012; Daub *et al.* 2013; Berg and Coop 2014; Field *et al.* 2016; Zeng *et al.* 2018; Edge and Coop 2019; Speidel *et al.* 2019; Lloyd-Jones *et al.* 2019). Applications using these methods, however, were often unknowingly biased by subtle patterns of population structure (Liu *et al.* 2018; Josephs *et al.* 2019; Rosenberg *et al.* 2019; Berg *et al.* 2019a; Sohail *et al.* 2019). Nonetheless, even considering the inflation of polygenic signals due to unrecognized population structure, mounting evidence over the last decade using a variety of methodologies, supports the polygenic adaptation model in a diversity of organisms, such as humans (Hancock *et al.* 2010a, b; Daub *et al.* 2013; Shi *et al.* 2016; Zeng *et al.* 2018; Gnecchi-Ruscone *et al.* 2018; Berg *et al.* 2019b), insects (Friedline *et al.* 2019), molluscs (Bernatchez *et al.* 2019), model plants (He *et al.* 2016), crops (Josephs *et al.* 2019; Wisser *et al.* 2019), and forest trees (Lind *et al.* 2017; De La Torre *et al.* 2019). However, there are still multiple open questions regarding the degree of polygenicity at adaptive traits, the distribution of effect sizes the involved loci, and how the genetic architecture changes under varying selective forces, especially for non-model species (Lind *et al.* 2018).

Following the expectations of the polygenic adaptation model, the heritability of complex traits is often associated with loci that are widespread across the whole genome, also including SNPs located in genes and pathways that do not show a clear functional connection to the trait of interest (Boyle *et al.* 2017). The omnigenic model, formulated by Boyle *et al*. (2017), provides a plausible hypothesis to explain this. The interconnection of gene regulatory networks implies that the vast majority of expressed genes likely influence the functions of core genes directly linked to fitness-related traits. Nevertheless, the study of polygenic adaptation at the pathway level is useful to identify gene sets of special relevance for adaptation (Daub *et al.* 2013). In particular, polygenic adaptation at pathway level has been proved especially useful to study non-model species, as reliance on selective sweep models often led to poor inferences (Hämälä *et al.* 2020). For example, Mayol *et al.* (2020) detected signatures of polygenic selection at the pathway level in English yew (*Taxus baccata* L.) *Taxus baccata* and identified negative selection as an important mechanism driving the pathway-level signal. Similarly, negative selection has been identified as a pervasive mechanism determining the polygenic architecture of fitness-related traits in humans (Zeng *et al.* 2018; O’Connor *et al.* 2019). In particular, negative selection has been proposed to favor polygenicity in complex traits by removing large-effect variants, because of their deleterious effects, while small-effect variants would remain unaffected; a process named ‘flattening’, as the genetic signal is ‘flattened’ relative to the underlying biology (O’Connor *et al.* 2019).

Despite theoretical advances and the development of new methods to study polygenic adaptation, the addressed questions remain constrained by the specific life-history traits of a few model species. Maritime pine (*Pinus pinaster* Ait.) is an ideal case study to investigate polygenic adaptation in an ecologically and economically important group of species, the forest trees: it is a long-lived plant inhabiting nearly undomesticated random-mating populations with high genetic diversity (González-Martínez *et al.* 2002; Jaramillo-Correa *et al.* 2015). It expanded from several isolated glacial refugia, and it is now distributed across the western Mediterranean Basin and the European Atlantic front in scattered populations under contrasting environments (Bucci *et al.* 2007). In addition, an artificial clonal propagation program in maritime pine allowed us to estimate precisely variance components and investigate selective forces driving trait evolution under contrasting environments. Specifically, we i) tested the hypothesis that most complex adaptive traits in a long-lived plant are polygenic, providing a first estimate of the degree polygenicity in a forest tree, and ii) that their genetic architecture is mostly driven by negative selection. We then iii) investigated how these patterns change with time and through environmental settings, which is of special relevance for long-lived organisms, such as forest trees.

## Materials and Methods

### Clonal common garden network (CLONAPIN)

We studied phenotypic variation in a clonal common garden network (CLONAPIN) composed of five sites covering the natural environmental range of maritime pine, from harsh Mediterranean climates to mild Atlantic ones. Common gardens comprise trees from 36 populations, sampled across the species natural distribution and covering the six previously identified gene pools for this species (Jaramillo-Correa *et al.* 2015; Supplemental Figure S1). Open-pollinated seeds were collected in natural stands, and germinated in a nursery; then one seedling per open-pollinated family was selected and vegetatively propagated by cuttings (following Majada *et al.* 2011). A total of 535 genotypes (clones) belonging to 35 populations were used to establish four of the clonal common gardens (three sites in Spain: Cabada, Cáceres and Madrid; and one in Portugal: Fundão; Table 1) with eight ramets per clone set in a randomized complete block design (*N*=4,273 trees). These clonal common gardens were planted in 2010. In 2011, a fifth common garden was established in Pierroton (France), comprising 443 clones from all 36 populations (*N*=3,434 trees). Common gardens in Cabada, Fundão and Pierroton are located in the Atlantic region, with high annual rainfall and mild temperatures. Common gardens in Cáceres and Madrid are located in continental areas under Mediterranean influence, with large seasonal temperature oscillations and a marked summer drought. In addition, clay soils in Cáceres hampered plant growth and diminished survival (Table 1).

**Table 1.** CLONAPIN common garden network (5 sites). Climatic data correspond to the mean of each parameter for the period 2005-2014 obtained from the EuMedClim database (Fréjaville and Benito Garzón 2018).

| Site | Country | Coordinates | Environment | Plantation<br>year | N trees<br>(clones) | Annual<br>precipitation<br>(mm) | Summer<br>precipitation<br>(mm) | Annual mean<br>temperature<br>(°C) | Annual<br>temperature<br>range (°C) | Soil type |
| --- | --- | --- | --- | --- | --- | --- | --- | --- | --- | --- |
| **Cabada** | Spain | 43°25'17" N<br>06°32'38" W | Iberian<br>Atlantic | 2010 | 4,272<br>(535) | 890 | 126 | 12.9 | 24.0 | Cambisol |
| **Fundão** | Portugal | 40°06'38" N<br>07°28'58" W |  | 2010 | 4,272<br>(535) | 1122 | 58 | 14.0 | 26.9 | Cambisol |
| **Pierroton** | France | 44°44'42" N<br>00°47'04" W | French<br>Atlantic | 2011 | 3,434<br>(443) | 933 | 199 | 13.8 | 26.7 | Arenosol |
| **Madrid** | Spain | 40°30'47" N<br>03°18'44" W | Mediterranean | 2010 | 4,272<br>(535) | 378 | 35 | 14.8 | 32.8 | Arenosol |
| **Cáceres** | Spain | 40°02'24" N<br>05°22'19" W |  | 2010 | 4,272<br>(535) | 374 | 21 | 16.7 | 32.6 | Fluvisol |

### Phenotypic evaluation

A total of 28 phenotypic trait-environment combinations were evaluated in this study. Assayed phenotypic traits were classified into five groups: survival, height, phenology-related, functional, and biotic-stress response (see Supplemental Table S1 for an exhaustive list of the measured traits). In brief, tree survival and height were evaluated in the five common gardens (including different years in Pierroton, with measures taken in 2013, 2015, and 2018). Phenology-related traits were evaluated in the Atlantic sites only (Cabada, Fundão and Pierroton), including different years of evaluation in Pierroton (2015 and 2017). In Cabada and Fundão, growth phenology was estimated through the presence of polycyclism (i.e. the ability for a plant to produce several flushes in the same growing season; (Girard *et al.* 2011)) and a phenology growth index (1):

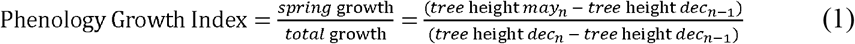

where *may* and *dec* correspond to the months May and December of the year n and the year n−1, respectively.

In Pierroton, phenology of bud burst was estimated through a scale ranging from 0 to 5 (0: bud with no winter elongation, 1: elongation of the bud, 2: emergence of brachyblasts, 3: brachyblasts begin to separate, 4: elongating needles, 5: total elongation of the needles) (see Hurel *et al.* 2019). The first Julian day at each stage (S1 to S5) was scored for each tree. Julian days were converted into accumulated degree-days (with base temperature 0°C) from the first day of the year, to take into account the between-year variability in temperature. The number of degree-days between stages 1 and 4 defines the duration of bud burst. Daily mean temperatures to calculate accumulated degree-days were downloaded from the nearest climatic station (located just a few hundred meters from the common garden, station 33122004 of the INRAE Agroclim database: https://www6.paca.inrae.fr/agroclim/Les-outils).

Functional traits, including nitrogen and carbon content and isotopic composition (δ^15^N and δ^13^C, respectively), as well as specific leaf area (SLA, a measure of leaf area per unit of dry mass), were evaluated in the common garden located in Fundão (Portugal). A bulk of five needles positioned 10 cm below the upper part of the shoot to avoid sampling bias (Warren *et al.* 2001) were sampled and prepared in a standard way for analysis (Brendel 2001). Determination of carbon content and isotopic composition was performed with a mass spectrometer at the University of Colorado isotope laboratory. Raw values were corrected by their position in the plate according to the standards, and this value was used for the subsequent analysis. SLA is an estimation of the compromise among light capture, CO_2_ assimilation, and the restrictions imposed by water loss through transpiration (Sefton *et al.* 2002). Low SLA suggests high leaf construction cost, and thus higher stress tolerance (Díaz *et al.* 2016). Thus, this key leaf trait is also associated with fitness components, such as tree survival (Greenwood *et al.* 2017). Given that there is a positive relationship between δ^13^C and water use efficiency (Farquhar and Richards 1984), δ^13^C has been widely used as a surrogate to study tree adaptation to water-limiting environments (e.g. Aranda *et al.* 2010; Walker *et al.* 2015). Similarly, δ^15^N is an indirect index related to the nitrogen cycle (Craine *et al.* 2015).

Assessment of biotic-stress response in a high number of trees is logistically complex. Therefore, it was evaluated only for a subset of clones (see Supplemental Table S1) in the Pierroton common garden (France). This common garden was selected because of the importance of the Landes region in maritime pine breeding for wood production. Biotic-stress response was evaluated based on susceptibility to two major pine pathogens, *Diplodia sapinea* and *Armillaria ostoyae*, as well as the incidence of the defoliator pest, *Thaumetopoea pityocampa* (pine processionary moth) (see Hurel *et al.* 2019 for details). *D. sapinea* causes several diseases in conifers, which may be exacerbated under climate change and compromise pine forest health (Desprez-Loustau *et al.* 2006). Susceptibility to *D. sapinea* was evaluated following controlled inoculation as the lesion extent around the inoculated site (hereafter referred as necrosis) and a scalar notation of needle discoloration (0: no discoloration, 2: some needles around the necrosis were discolored, and 3: all needles around the necrosis were discolored). *A. ostoyae* is a conifer root pathogen causing growth cessation and eventually death (Heinzelmann *et al.* 2019). To evaluate the incidence of this pathogen, *A. ostoyae* mycelium culture was prepared in plastic jars. The level of humidity observed in the plastic jar was visually scored as dry, medium or humid. Susceptibility to *A. ostoyae* was assessed after controlled inoculation as the lesion length in the sapwood (i.e. wood browning, hereafter also referred as necrosis). We accounted for the potential influence of variation in humidity on wood browning by including the level of humidity in the jar as a covariate for *A. ostoyae* susceptibility analysis (see below). Finally, the pine processionary moth is an insect that rapidly defoliates pines leading to forest decline (Jacquet *et al.* 2013). The presence or absence of pine processionary moth nests in the tree crowns was assessed in March 2018.

### DNA extraction and SNP genotyping

Needles were collected from one ramet per clone in the Cabada common garden (*N*=523). Genomic DNA was extracted using the Invisorb® DNA Plant HTS 96 Kit/C kit (Invitek GmbH, Berlin, Germany). A 9k *Illumina Infinium* SNP array developed by Plomion *et al.* (2016b) was used for genotyping. This array was constructed using previously identified and newly *in silico-*developed SNPs, either from randomly screened EST sequences or specifically detected at candidate genes for adaptation to biotic and abiotic factors (see Plomion *et al.* 2016b for further details). Genotyped SNPs covered all 12 chromosomes of *P. pinaster* according to previous linkage mapping (Plomion *et al.* 2016b). For this study, 6,100 SNPs were finally retained following standard filtering (GenTrain score > 0.35, GenCall50 score > 0.15 and Call frequency > 0.85) and removal of SNPs with uncertain clustering patterns (visual inspection using *GenomeStudio v. 2.0*). Individuals with more than 15% missing data were also removed. This resulted in 5,165 polymorphic SNPs that were included in the estimation of molecular population differentiation (*F_ST_*) and the polygenic association study.

### Quantitative genetics analysis

Genetic components of the phenotypic variance were estimated using Generalized Linear Mixed-Effects Models (GLMM) fitted in a Bayesian framework using Markov chain Monte Carlo (MCMC) methods. The model, described in equation (2), was implemented for those phenotypic traits evaluated at multiple sites of the CLONAPIN common garden network (see Supplemental Table S1). To estimate the genetic control of the genotype-by-environment (G×E) interactions, the model described in equation (3) was fitted for those traits measured at all sites of the CLONAPIN common garden network (i.e. height and survival).

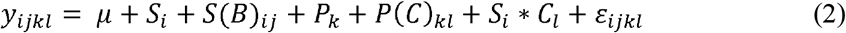

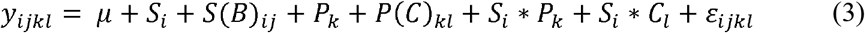

where, for a given trait *y*, *μ* denotes the overall phenotypic mean, *S_i_* refers to the fixed effect of site *i, B_j_* represents the random effect of experimental block *j* nested within site *i*, *P_k_* is the random effect of population *k*, *C* denotes the random effect of clone *l* nested within population *k*, and *ε* is the residual effect.

Simplified models with or without covariates represented by equations (4) and (5) were implemented for phenotypic traits measured in just one site of the CLONAPIN common garden network (see Supplemental Table S1).

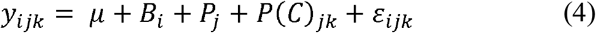

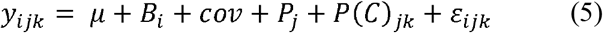

Where, for a given trait *y*, *μ* denotes the overall phenotypic mean, *B_i_* represents the fixed effect of experimental block *i*, *P_j_* is the random effect of population *j*, *C* denotes the random effect of clone *k* nested within population *j*, and *ε* is the residual effect. In the model represented by equation (5), *cov* represents a covariate implemented when modeling the presence of pine processionary moth nests (i.e. an estimate of tree height) and necrosis caused by *A. ostoyae* (i.e., level of humidity in the experimental jar).

All models were fitted with the R package *MCMCglmm* (Hadfield 2010). Phenotypic traits showing Gaussian distributions where modeled using the identity link function, while phenotypic traits exhibiting a binomial distribution (survival, polycyclism, *D. sapinea* needle discoloration, and presence or absence of pine processionary moth) were modeled either with *logit* or *probit* link functions (see Supplemental Table S2 for an exhaustive list of model parameter specifications). Multivariate-normal prior distributions with mean centered at zero and large variance matrices (10^8^) were used for fixed effects. For ordinal traits, a Gelman prior for the variance of fixed effects was set, as suggested by Gelman et al. (2008). Inverse Wishart non-informative priors were used for the variances and covariances of random effects, with the matrix parameter *V* set to 1, and a parameter *n* set to 0.002 (Hadfield 2010). Parameter expanded priors were used to improve the chain convergence and mixing, as suggested by Gelman (2006) for models with near-zero variance components. Priors with a larger degree of belief parameter (*n* set to 1), specifying that a large proportion of the variation is under genetic control (as suggested by Wilson *et al.* 2010) did not change the results (data not shown). Models were run for at least 550,000 iterations, including a burn-in of 50,000 iterations and a thinning interval of 100 iterations. Four chains per model were run to test for parameter convergence. The potential scale reduction factor (psrf) was consistently below 1.02 for all the models (Supplemental Table S2) (Gelman and Rubin 1992).

Variance components were then used to compute broad-sense heritability (*H^2^*) as (6):

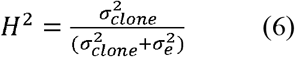

where 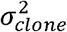 is the variance among clones within populations and 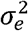 the residual variance. For estimating broad-sense heritability for traits following a binomial distribution, we included an extra term in the denominator (+ π^2^/3) to account for implicit *logit* link function variance; similarly, we added one to the denominator to account for the *probit* link function (Nakagawa and Schielzeth 2010).

The GLMMs described above were used to estimate genetic values using Best Linear Unbiased Predictors (BLUPs) (Henderson 1973; Robinson 1991). The genetic value of each clone was defined as the population BLUP plus the clone BLUP. BLUPs for G×E, were obtained from equation (3) and calculated following equation (7).

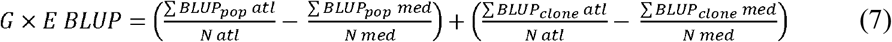

where *BLUP_pop_ atl* is the population BLUP in sites under Atlantic climate (Cabada, Fundão, and Pierroton), *BLUP_pop_ med* the population BLUP in sites under Mediterranean climate (Cáceres and Madrid), *BLUP_clone_ atl* the clone BLUP in sites under Atlantic climate*, BLUP_clone_ med,* the clone BLUP in sites under Mediterranean climate, *N atl* the number of sites under Atlantic climate, and *N med* the number of sites under Mediterranean climate.

Parameter estimates from quantitative genetics analyses are presented as the mode of the posterior distribution; 95% credible intervals were computed as the highest density region of each posterior parameter distribution.

### Q_ST_-F_ST_ comparison

Molecular population differentiation (*F*_ST_) was estimated according to Weir and Cockerham (1984) using the 5,165 SNPs from the *Illumina Infinium* SNP array and the diveRsity R package (Keenan *et al.* 2013). The 95% confidence interval of the global *F_ST_* estimate was computed by bootstrapping across loci (1,000 bootstrap iterations). Quantitative genetic differentiation among populations was calculated following Spitze (1993) using the variance components estimated from the previously described models (equations 2-5):

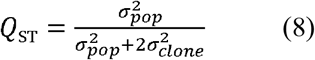

where 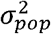 is the variance among populations, and 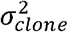 is the variance among clones within populations. Quantitative (*Q*_ST_) and molecular (*F*_ST_) genetic differentiation among populations were considered significantly different when *Q*_ST_ and *F*_ST_ posterior distributions had non-overlapping 95% confidence intervals.

### Polygenicity across traits, years and environments

Polygenicity was evaluated as the proportion of SNPs with non-zero effects on phenotypic traits. First, we conducted posterior inference via model averaging and subset selection (VSR), as implemented in piMASS software (Guan and Stephens 2011). This method allows to identify combinations of SNPs likely affecting a phenotype and to estimate the proportion of trait variance explained by the SNPs in the data set. Hereafter, we referred to this quantity as the genetic explained variance (*GEV*), which, in this study, represents the BLUP variance explained by SNP additive effects. Second, we used the Bayesian mixed linear model (MLM) framework developed by Zeng *et al.* (2018) as implemented in CGTB 2.0 software. This last model simultaneously estimates: i) SNP-based heritability (considering the SNPs with non-zero effects on the trait), hereafter referred as *GEV*, analogously to VSR estimates, ii) polygenicity (as defined above), and iii) the relationship between SNP effect-size and minor allele frequency (*S*, a common indicative of negative selection). When negative selection is operating, *S* is expected to be negative, as most new mutations are deleterious and high-effect SNPs are kept at low frequencies. Estimates with 95% credible intervals of parameter posterior distributions not overlapping zero were considered as significant. Prior to these analyses, neutral population genetic structure was accounted for by running linear models relating the genetic values for each trait (with site and block effects removed) to the admixture coefficients for each clone (*Q*-scores) obtained using a STRUCTURE run for *K*=6 based on neutral markers (see Jaramillo-Correa *et al.* 2015 for further details). From this linear model, we extracted the normalized residuals for each trait, as recommended in piMASS manual.

Analyses were run separately for different traits, years, and environments (see Supplemental Table S1). VSR models were run for 2,000,000 iterations with a burn-in of 100,000 iterations and a thinning interval of 100. After several preliminary runs, the maximum number of SNPs included in VSR models was fixed to 2,000 (i.e. maximum allowed polygenicity of ~40%). MLM models were run for 500,100 iterations, including a burn-in of 100 iterations, and a thinning interval of 10 iterations. Parameter estimates from both VSRs and MLMs were presented as the median of the posterior distribution, instead of the mode, for better handling of bimodal distributions (Supplemental Figure S2). The 95% credible intervals were computed as the highest density region of the posterior parameter distribution, as above.

### Annotation and gene function enrichment at pathway level

The transcripts containing the 5,165 polymorphic SNPs were downloaded from SustainPine v.3.0 database (Canales *et al.* 2014). DNA sequences were translated with BioEdit v. 7.2.6 (Hall 1999) and submitted to BlastKOALA (Kanehisa *et al.* 2016) for annotation and functional characterization using InterPro annotations, GO terms, and KEGG pathway identification. Annotations were compared with those available at SustainPine, and conflicting cases were examined individually by privileging similarity to genes correctly identified in other conifers or forest trees. Contigs with no clear annotations (e.g. hypothetical or unknown proteins, or unsolved conflicting annotations) were removed from the database. For the retained contigs, the top-two KEGG terms were used for assignation to one or more specific metabolic pathways/modules based on KEGG orthology. Genes for which no hit with KEGG database was found, were assigned to metabolic pathways/modules based on the InterPro annotation. We privileged metabolic pathways/modules that could be unequivocally assigned to a given phenotypic response (e.g. circadian rhythm to bud phenology or pathogen interaction to biotic stress response) or linked to various stress responses (e.g. DNA recombination and repair, ubiquitin system or transcription factor machinery to survival and biotic stress response). In total, seventeen pathways/modules were retained containing a total of 628 (19.7% out of 3,194) genes, with 1,233 polymorphic SNPs (Supplemental Table S3).

For enrichment tests using polysel (Daub *et al.* 2013), the seventeen pathways/modules were defined as gene sets. First, we computed two statistics at the gene level (i.e. objStat in polysel) based on the per-SNP estimates obtained from the VSR implemented in piMASS: the maximum, over all SNPs included in a gene, of the Rao-Backwellized posterior probability of inclusion *maxpostrb*, and the maximum of the absolute value of Rao-Backwellized effect size *maxabsbetarb*. To account for a weak correlation of these statistics with the number of SNPs per gene, we used the AssignBins and RescaleBins functions in polysel, which automatically assigns gene scores (objStat) into bins defined from the number of SNPs per gene. We then rescaled scores within bins and computed the sum(objStat) of each statistic over all genes per gene set. Since the sum(objStat) for random gene sets (sizes n = 10, 50, 250 genes) was not normally distributed, we built empirical null distributions by randomly sampling gene sets of the same size as the sets to be tested. Then, we performed one-sided tests evaluating whether the observed sum(objStat) was smaller than the 5^th^ or larger than the 95^th^ percentile of the sum(obStat) null distribution. Higher-tail significant results for *maxpostrb* indicate gene sets enriched with higher overall probability of being selected during the VSR procedure implemented in piMASS. Higher-tail significant results for *maxabsbetarb* points to gene sets enriched with higher overall SNP effect-sizes. Contrarily, lower-tail significant results for both statistics suggests conserved gene sets, containing genes with smaller overall probability of inclusion or SNP effect-size estimates. We report *p*-values based on this comparison, as well as *q*-values from a False Discovery Rate (FDR) approach implemented in the R package *qvalue* (R Core Team 2019). The level of connection between gene sets was weak with only four genes associated with more than one gene set (633 gene – gene set combinations for 628 genes). For this reason, we did not assess enrichment for pruned gene sets (see Daub *et al.* 2013).

## Results

### Broad-sense heritability and genetic differentiation among populations

All traits had low to moderate estimates of broad-sense heritability (Supplemental Table S1), with the exception of nitrogen and carbon amount that did not show genetic variation. Consequently, polygenic association methods failed to converge for these two traits and they were excluded from further analyses. *H^2^* ranged from 0.32 for bud burst measured in 2015 to zero for survival in the French Atlantic environment in 2013. Interestingly, survival showed significant estimates of *H^2^* only in the sites under (harsher) Mediterranean climate. The highest *H^2^* estimates were observed for phenology-related traits followed by tree height. *H^2^* for a given trait varied across environments (e.g. height, survival and phenology-related traits) but showed little variation across years (Supplemental Table S1).

The global *F*_ST_ was 0.112 (95% confidence interval: 0.090 - 0.141). All groups of phenotypic traits, excepting survival, had at least one trait with statistically higher *Q*_ST_ than *F*_ST_ (Supplemental Table S1). The highest *Q*_ST_ was obtained for susceptibility to *D. sapinea* infection measured as necrosis length, followed by δ^13^C and tree height, which also showed similar *Q*_ST_ values across environments and tree ages (Figure 1).

**Figure 1.**
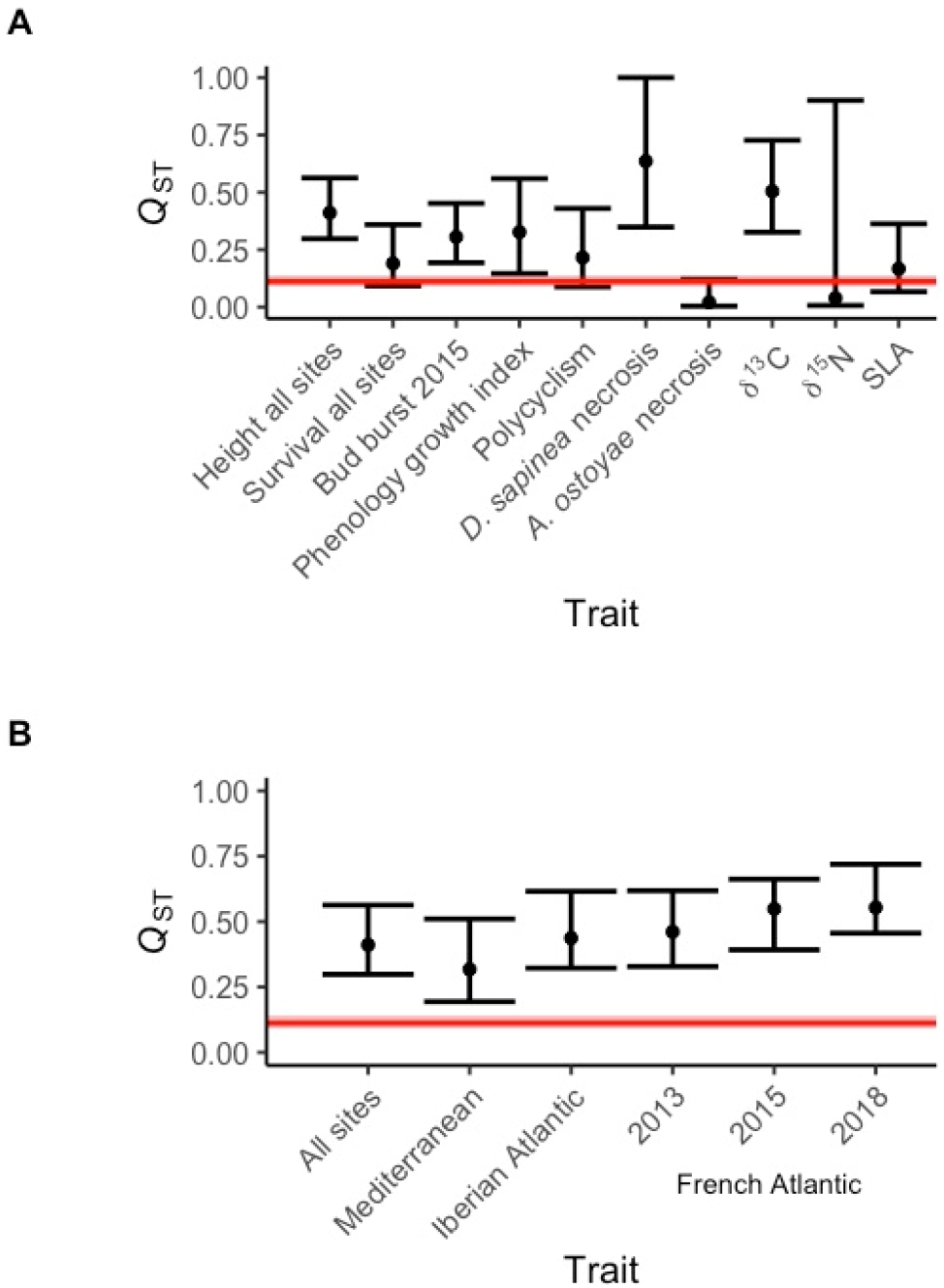
Comparison of *Q*_ST_ and *F*_ST_ estimates across traits, environments and years. A) *Q*_ST_ for a selection of traits belonging to five categories: survival, height, phenology-related traits, functional traits and biotic-stress response (see Supplemental Table S1 for all traits). B) *Q*_ST_ for height estimated in three different environments: Mediterranean, Iberian Atlantic, and French Atlantic, and a global *Q*_ST_ for the three environments together. In the French Atlantic common garden, height was measured in three different years: 2013, 2015 and 2018. Global *F*_ST_ estimate is presented by a red line surrounded by the 95% confidence intervals computed by bootstrapping.

### Genetic architecture (polygenicity) of adaptive traits

Polygenicity estimates were consistent between the VSR and MLM methods (Supplemental Table S4). Both methods showed substantial polygenic control for most of the phenotypic traits, with an average of 6% (0-27%) of the genotyped SNPs having non-zero effects. Significant polygenicity was found in all five trait categories for at least one trait (Figures 2 and 3; Supplemental Table S4). Polygenicity for height was stable across environments and years, when measured multiple times under the same environment (i.e. in the French Atlantic common garden) (Figure 3). Along the same line, polygenicity for phenology-related traits and tree survival also remained stable across environments, although 95% credible intervals overlapped zero in some cases. The low polygenicity values observed for survival in the French Atlantic common garden are probably a consequence of the low levels of phenotypic variability in this site, with almost no mortality (97.12% of planted trees were alive at the evaluation time, Supplemental Table S1). Polygenicity was heterogeneous for biotic-stress response and functional traits (Figure 2). For instance, susceptibility to *D. sapinea* was more polygenic than to *A. ostoyae* or than incidence of pine processionary moth. For functional traits, SLA and δ^15^N showed the highest levels of polygenicity, while δ^13^C showed a considerably lower proportion of SNPs with non-zero effects.

**Figure 2.**
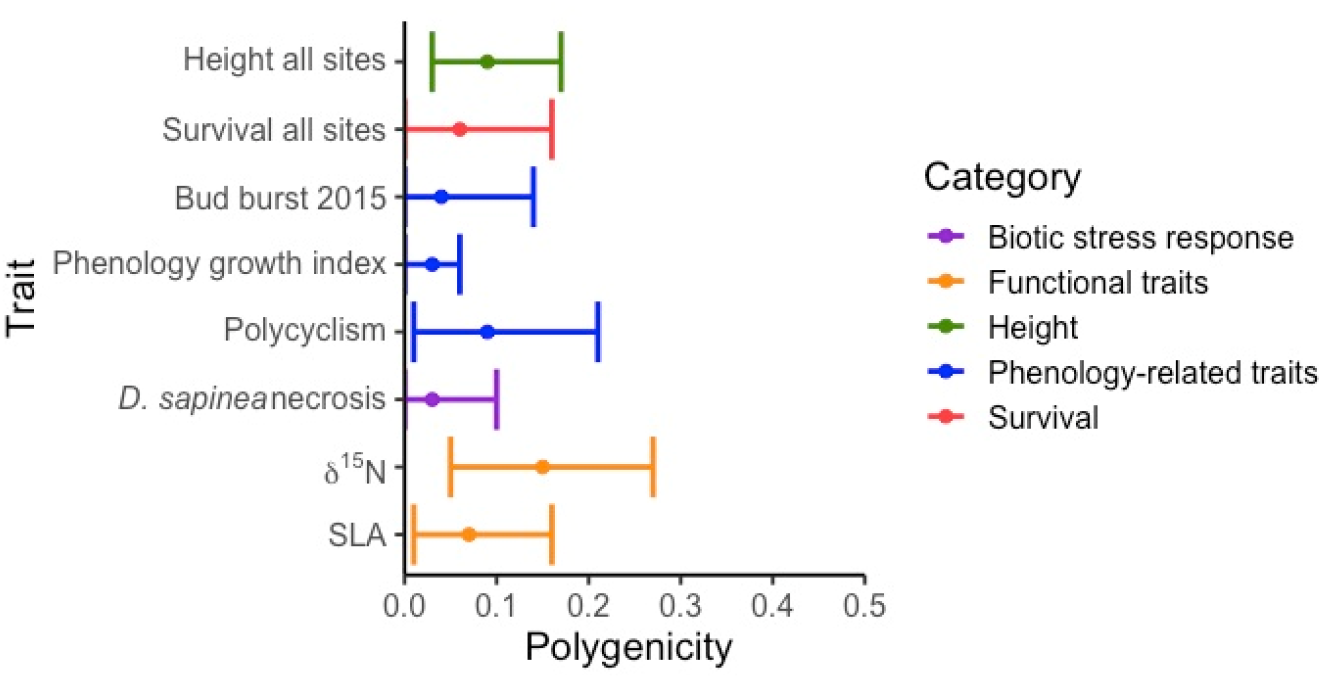
Polygenicity estimated from Bayesian mixed linear models (MLMs) for a selection of traits (see Supplemental Table S4 for all traits). Polygenicity was estimated as the proportion of non-zero size-effect SNPs. Posterior median and 95% credible intervals are presented.

**Figure 3.**
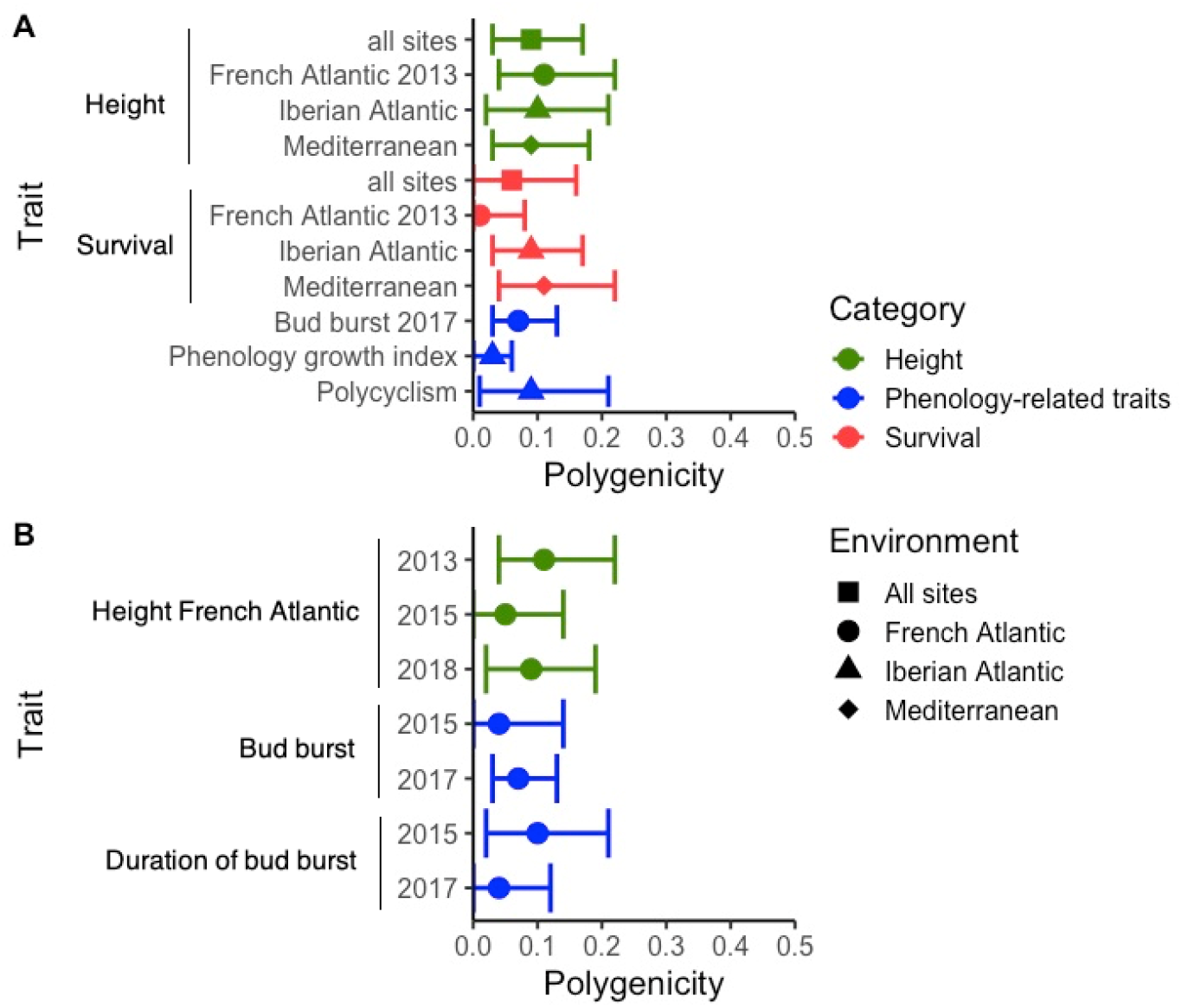
Polygenicity estimated from Bayesian mixed linear models (MLMs) across environments and years. A) Variation of polygenicity across environments. B) Temporal variation of polygenicity. Polygenicity was estimated as the proportion of non-zero size-effect SNPs. Posterior median and 95% credible intervals are presented.

In addition, *GEV* was consistent between methods, although VSR tended to render higher values (Supplemental Table S4). On average *GEV* was 0.38 across traits, with a minimum of 0.018 for survival in the French Atlantic environment in 2018, and a maximum of 0.99 for *D. sapinea* necrosis. *GEV* estimated with the VSR method for the G×E component on tree height (considering Atlantic versus Mediterranean environments) was low but significant (median = 0.238, 95% credible interval = 0.043 - 0.409), indicating some SNPs with significant effects on growth plasticity. However, this result could not be confirmed with the MLM method. Moreover, *GEV* for the G×E component on tree survival was not significant with any model.

Polygenicity and *GEV* were positively and consistently correlated for both VSR and MLM models (Figure 4). This positive correlation suggested that SNP-based heritability is mainly determined by genetic variants with similarly small effects, and that differences in polygenicity across traits are mostly accounting for differences in explained genetic variance, rather than the distribution of SNP effect-size (Supplemental Figures S2 and S3).

**Figure 4.**
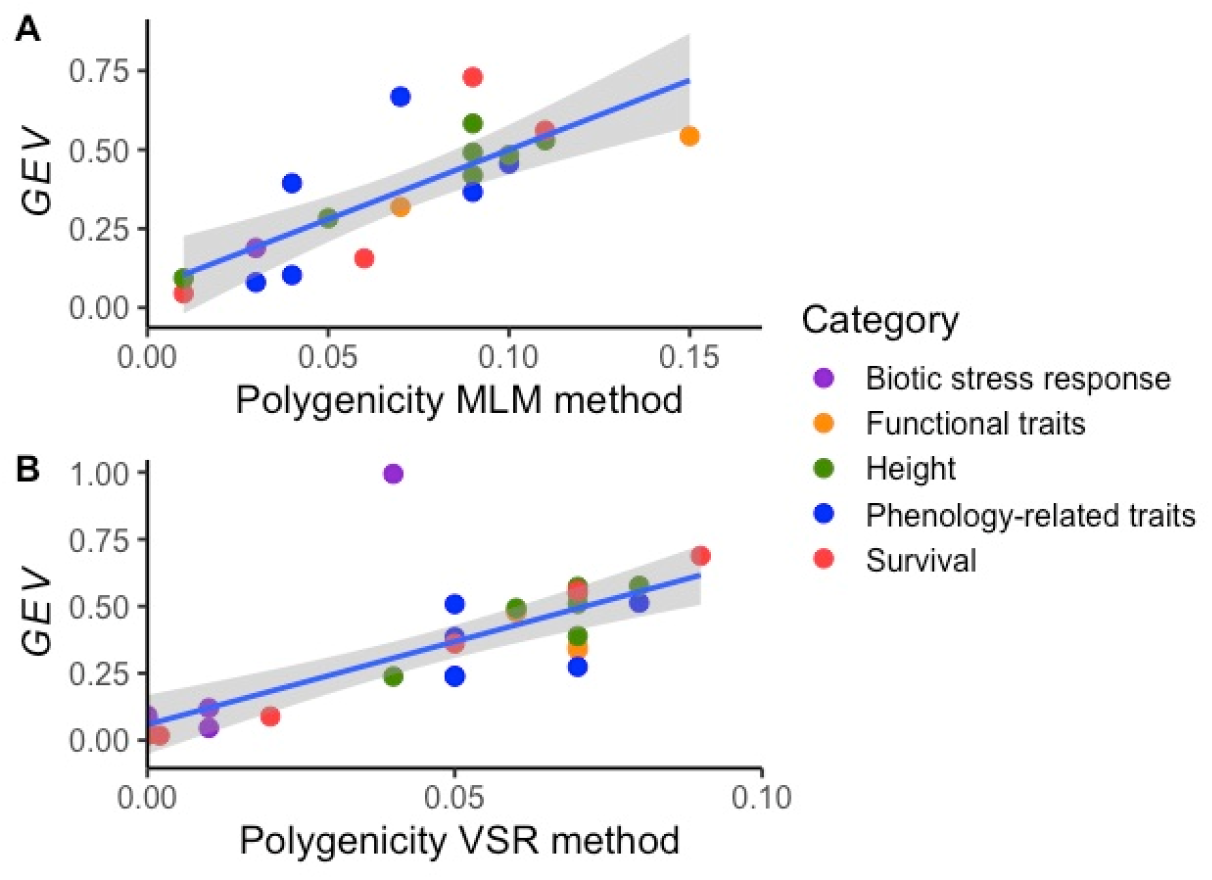
Correlation between polygenicity (proportion of non-zero size-effect SNPs) and *GEV* (explained genetic variance). A) MLM method implemented in CGTB software. B) VSR method implemented in piMASS software. Each point represents the posterior median.

### Evidence of negative selection

The correlation between SNP effect-size and minor allele frequency (MAF), *S*, was used to identify the type and mode of natural selection acting upon phenotypic traits. Out of the 28 assayed traits, we were able to estimate *S* through the MLM method for 19 of them. Estimates ranged from −1.68 (bud burst in 2017) to 0.55 (tree survival in French Atlantic environment), but only seven traits from four out of five trait categories (survival, height, phenology-related, and functional traits) were significant (Figure 5). No significant effect was observed for any trait belonging to the biotic-stress response category. Remarkably, all seven significant estimates of *S* were negative (ranging from −1.68 for bud burst in 2017 to −0.99 for survival in the Iberian Atlantic environment).

**Figure 5.**
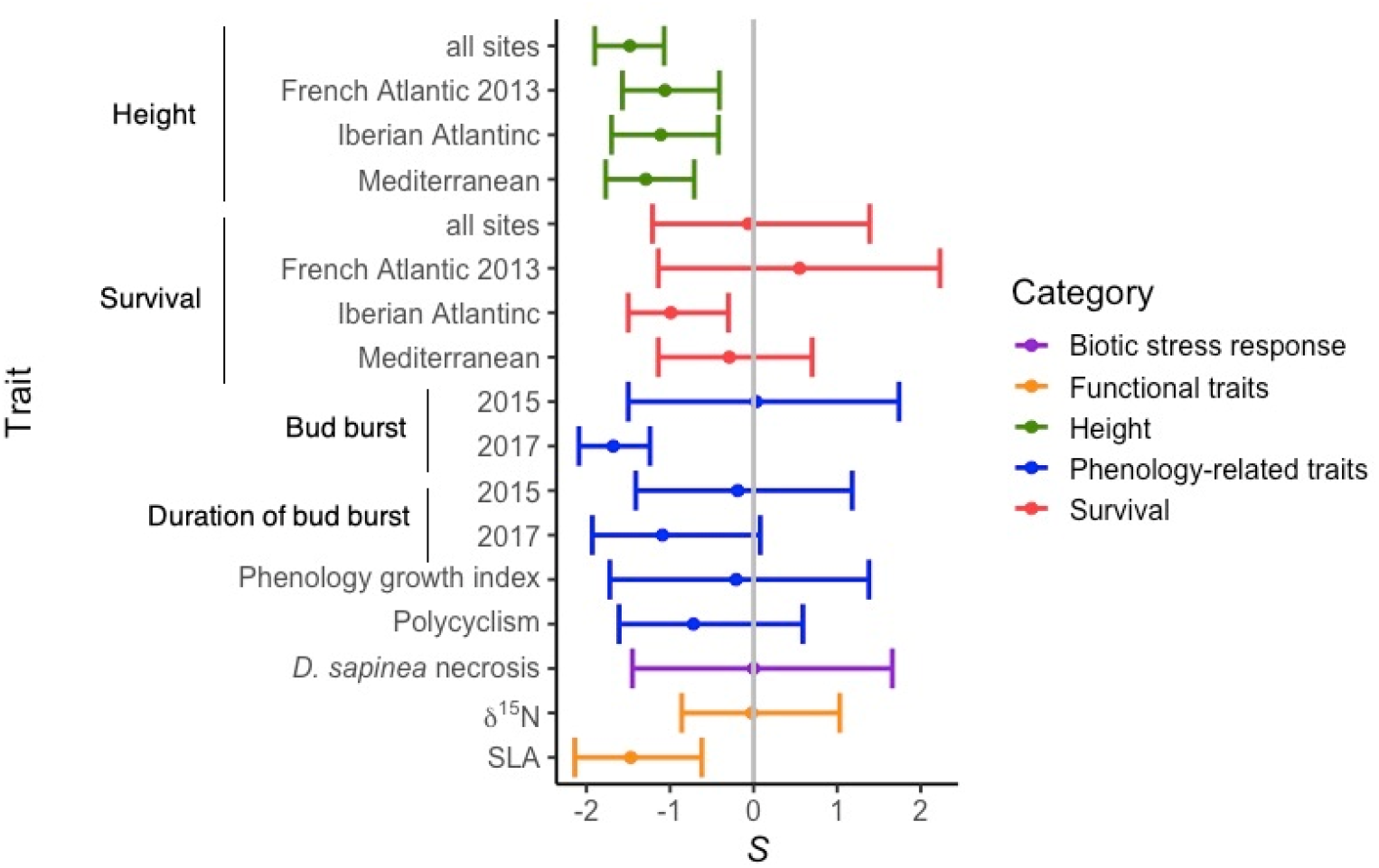
Correlation between SNP effect-size and Minor Allele Frequency (MAF). The coefficient of correlation between SNP effect-size and MAF (*S*) was estimated through the MLM method. The posterior distribution of *S* (median and 95% credible intervals) are presented.

Estimates of *S* for height were consistent across years and environments. However, *S* estimated for tree survival was only significant in the Iberian Atlantic environment. For phenology-related traits, *S* was significant only for bud burst measured in 2017 (Figure 5). These results contrast with the consistent level of polygenicity for all these traits across years and environments. Interestingly, our results suggest a stable polygenic architecture, but an environment and year-dependent impact of negative selection at some traits.

### Gene function enrichment at pathway level

Tests for gene function enrichment at the pathway level provided significant results for survival in the Iberian Atlantic environment, phenology-related and biotic-stress response traits, and height in the French Atlantic and Mediterranean environments. Genes coding for *transcription factors* showed higher probability of being included in the VSR models (*maxpostprb)* and higher estimated SNP effect-sizes (*maxabsbetarb)* for survival in the Iberian Atlantic environment (Table 2). Two gene sets associated to bud burst in 2015 showed signals of polygenic selection: *monolignol biosynthesis*, which had high overall values of both *maxpostprb* and *maxabsbetarb*, and *glycan metabolism*, which showed low overall *maxabsbetarb* estimates (Table 2). Furthermore, phenology growth index was associated with enrichment for genes related to *cell growth and death*, *DNA recombination and repair* and *UV response*, which mostly have low *maxabsbetarb* values (Table 2). *D. sapinea* susceptibility was associated with enrichment of genes, with high overall *maxabsbetarb* and *maxpostprb,* in the *ubiquitin system* for *D. sapinea* necrosis, and in the *signal transduction* and *flavonoid biosynthesis* gene sets for *D. sapinea* needle discoloration (Table 2). Interestingly, height was enriched for genes from different pathways when measured in contrasting environments. For instance, in the French Atlantic environment genes coding for *transcription factors* showed high *maxabsbetarb* and *maxpostprb*, while genes within the *cytoskeleton* pathway showed overall low *maxabsbetarb* values in the Mediterranean environment.

**Table 2.** Gene sets with gene function enrichment at pathway/module level. Two statistics obtained from the VSR method were tested: the maximum of any SNP per gene of the Rao-Backwellized posterior probability of inclusion (*maxpostprb*) and the maximum of any SNP per gene of the absolute value of the Rao-Backwellized effect-size (*maxabsbetarb*). Sign of enrichment refers to two-tailed null hypothesis testing.

| **Trait** | **Environment** | **Gene set** | **Statistic tested** | **Sign of enrichment** | **p-value** | **q-value (<0.10)** |
| --- | --- | --- | --- | --- | --- | --- |
| Height | French Atlantic | *Transcription factor* | *maxabsbetarb* | Higher | 0.003 | 0.05 |
|  |  |  | *maxpostprb* | Higher | 0.004 | 0.07 |
|  | Mediterranean | *Cytoskeleton* | *maxabsbetarb* | Lower | 0.003 | 0.06 |
| Survival | Iberian Atlantic | *Transcription factor* | *maxabsbetarb* | Higher | 0.001 | 0.01 |
|  |  |  | *maxpostprb* | Higher | <0.001 | 0.005 |
| Bud burst 2015 | French Atlantic | *Monolignol biosynthesis* | *maxabsbetarb* | Higher | 0.003 | 0.05 |
|  |  | *Monolignol biosynthesis* | *maxpostprb* | Higher | 0.005 | 0.08 |
|  |  | *Glycan metabolism* | *maxabsbetarb* | Lower | 0.040 | 0.09 |
| Phenology growth index | Iberian Atlantic | *Cell growth and death* | *maxabsbetarb* | Lower | 0.010 | 0.03 |
|  |  | *DNA recomb and repair* | *maxabsbetarb* | Lower | 0.008 | 0.03 |
|  |  | *UV response* | *maxabsbetarb* | Lower | 0.005 | 0.03 |
| *D. sapinea* necrosis | French Atlantic | *Ubiquitin system* | *maxabsbetarb* | Higher | 0.002 | 0.04 |
|  |  |  | *maxpostprb* | Higher | 0.003 | 0.06 |
| *D. sapinea* discoloration | French Atlantic | *Signal transduction* | *maxabsbetarb* | Higher | 0.004 | 0.08 |
|  |  |  | *maxpostprb* | Higher | 0.003 | 0.06 |
|  |  | *Flavonoid biosynthesis* | *maxpostprb* | Higher | 0.007 | 0.07 |

## Discussion

Unraveling the genetic architecture of adaptive traits is challenging because of the difficulty to identify variants with small effect-sizes using GWAS. Here, we addressed this challenge obtaining precise phenotypic information (over 12,500 trees were evaluated) for an extensive number of fitness-related traits measured on clonal replicates. Specifically, we tested if a high proportion of the genetic variance of fitness-related traits in a long-lived forest tree (maritime pine) can be explained by a large number of small size-effect variants, in line with the polygenic adaptation model. We also tested whether negative selection is pervasive for such polygenic traits. Our results showed patterns of local adaptation for most of the analyzed traits, highlighting its relationship with fitness, and also revealed a high and remarkably stable degree of polygenicity, across traits, years, and environments. Moreover, using two complementary multilocus approaches we accounted for a considerable proportion of the heritability estimated for these highly polygenic traits, and identified negative selection as a key driver of local adaptation.

### Evidence of local adaptation in maritime pine

All phenotypic categories presented significant within-population genetic variation (i.e. broad-sense heritability), and were consequently susceptible to respond to natural selection (Visscher *et al.* 2008). Estimates of heritability were consistent with previous results for these traits in forest trees (reviewed by Lind *et al.* 2018). In addition, our results were consistent with adaptive differentiation (*Q*_ST_ > *F*_ST_) for 11 out of 26 analyzed traits, involving four out of the five trait categories (no evidence for survival traits). These results are in accordance with reports of pervasive local adaptation in forest trees (Savolainen *et al.* 2007, 2013; Alberto *et al.* 2013; Lind *et al.* 2018).

The stability of *Q*_ST_ estimates for height across environments and years highlights the strength of directional selection for height in this species; a trait that can thus be used for the delimitation of conservation and management units (Rodríguez-Quilón *et al.* 2016). Contrarily, phenology-related traits showed contrasting estimates of *Q*_ST_ depending on the environment and year of measurement. This result highlights that the evolutionary forces driving population genetic differences in phenology-related traits are environmentally and temporally-dependent, which can slow-down attaining phenotypic optima under rapidly changing climates. Polygenic adaptation could be specially relevant for these traits because it can produce rapid phenotypic changes, as it only requires small adjustments in allele frequencies in the contributing loci rather than selective sweeps on new mutations (Jain and Stephan 2017; Dayan *et al.* 2019; Wisser *et al.* 2019).

Unexpectedly, survival, a trait directly related with a component of fitness (i.e. viability), did not show evidence of local adaptation in maritime pine. The low levels of phenotypic variability observed for survival in this study may explain these results. Future studies should focus on quantitative evaluations of survival (e.g. adding a time-frame, such as time until death or order of dead trees) to better gather the complexity of this trait, and be able to discern genetic differences among populations. The strong selective pressure in the Mediterranean region exacerbated genetic differences in survival among clones and resulted in slightly higher estimates of heritability (similarly to Gaspar *et al.* 2013). Additionally, we observed significant phenotypic plasticity for height and survival, the two traits measured in all five experimental sites. While our results hinted a heritable component for plasticity, this question still deserves further investigation to elucidate the importance of phenotypic plasticity in the adaptive response of maritime pine to changing environmental conditions (Alía *et al.* 2014; Vizcaíno-Palomar *et al.* 2019).

Two traits in particular had remarkably high levels of adaptive genetic differentiation among populations, δ^13^C and *D. sapinea* necrosis (Figure 1), but genetic variation within populations was low, compromising their adaptive potential. These traits deserve special attention because of the implication of water-use efficiency in drought resistance (reviewed by Plomion *et al.* 2016a) and the new pathogenic outbreaks of *D. sapinea* expected on maritime pine plantations fostered by climate change (Fabre *et al.* 2011; Brodde *et al.* 2019). In contrast to our findings, a lack of adaptive genetic differentiation for δ^13^C was previously reported for maritime pine by Lamy *et al.* (2011), as well as for broad-leaved trees (Torres-Ruiz *et al.* 2019). Although this disagreement may be influenced by the much larger number of populations we analyzed (see Whitlock and Guillaume 2009) as compared to Lamy *et al.* (2011), we cannot rule out discrepancies due to the estimation of total genetic variance in our study (i.e. based on clones), instead of additive genetic variance. Nevertheless, non-additive genetic effects in maritime pine traits related to drought resistance have been reported to be of little importance (Gaspar *et al.* 2013), and they should not have affected our estimates.

### Genetic architecture (polygenicity) of fitness-related traits

Most traits assessed had a considerable degree of polygenicity, ranging between 4-15%, which is on the same order of magnitude as for humans (Zeng *et al.* 2018). Polygenicity was relatively similar across all analyzed traits and therefore did not depend on the level of genetic control, as estimated by heritability through quantitative genetic analysis. Mei *et al*. (2018) observations in humans predicted different genetic architectures as a function of genome size. Surprisingly, although the maritime pine genome is more than seven times larger than that of humans (De La Torre *et al.* 2014), we found similar estimates of polygenicity between both species. The distributions of SNP effect-sizes showed that hundreds of SNPs with near-zero effect-size contributed together to shape phenotypic differences among clones. This highly polygenic architecture could be explained by the omnigenic model (Boyle *et al.* 2017). Indeed, as in humans, we expect high biological complexity and interconnectivity of gene expression networks in forest trees, resulting in the association of virtually all expressed genes in relevant tissues with the observed phenotypes (Wray *et al.* 2018). However, this explanation would not account for the lack of high effect-size SNPs in our data set composed mostly of SNPs from candidate genes (see below).

The implementation of polygenic adaptation studies outside of humans is slowly emerging (Csilléry *et al.* 2014; He *et al.* 2016; Lind *et al.* 2017; Barghi *et al.* 2019; Friedline *et al.* 2019; Wisser *et al.* 2019), providing increased evidence that polygenic adaptation in complex traits may be pervasive (Sella and Barton 2019). As a result, new evolutionary questions relevant for different organisms are arising. For instance, in forest trees, for which local adaptation is frequently observed (Savolainen *et al.* 2007, 2013; Alberto *et al.* 2013; this study), the contribution of alleles with small effect-size and selection coefficients (and therefore more prompted to be swamped by gene flow) to shaping local adaptation is a question that remains open (Yeaman 2015). Another fundamental question, in particular for conifers, is the role of genetic redundancy. It has been suggested that genetic redundancy favors polygenic adaptation and speed up the achievement of phenotypic optima through multiple genetic pathways leading to similar phenotypes (Höllinger *et al.* 2019; Barghi *et al.* 2019). Unraveling this relationship in conifers, whose genomes are characterized by a high number of paralogs (Diaz-Sala *et al.* 2013), may shed new light about how rapidly these taxa can adapt to environmental changes. Moreover, the influence of genome size in the genetic architecture of fitness-related traits, as well as the relationship between heritability and polygenicity, deserve further investigation including a better coverage of conifer genomes, as well as improved knowledge of non-coding regions (Mackay *et al.* 2012).

Recent studies in human height (a classic example of polygenic adaption) have suggested that detecting polygenicity may be affected by subtle biases in GWAS caused by population structure (Berg *et al.* 2019a; Sohail *et al.* 2019). In our study, the clonal common garden network allowed separating the genetic and the environmental effect on phenotypes to identify which traits are contributing to adaptation. In addition, we corrected the BLUPs estimates for the effect of neutral population genetic structure. In this sense, our work highlights the potential of combining precise estimation of the genetic effect on phenotypes with multi-locus genotype-phenotype association models to elucidate the mechanisms that allow the maintenance of genetic variation in adaptive traits, especially in species with complex demographic history. Undoubtedly, next steps to decipher polygenic adaptation in species with varied life-history traits should implement upcoming polygenic association methods that directly correct for population stratification (e.g. Josephs *et al.* 2019).

### Performance of polygenic adaptation approaches (VSR and MLM)

We evaluated the performance of polygenic approaches (VSR and MLM) through the comparison of SNP-based genetic variance estimates, *GEV.* Despite some slight differences, notably for biotic-stress response traits that were limited by low sample sizes, both methods were robust and provided consistent estimations. The large proportion of the genetic variance explained by SNP-based models, usually higher than 50%, suggests that, by adopting a polygenic analytical model, we were able to account for a significant part of the heritability inferred through pedigree-based analysis, even when using a modest number of SNPs. It is worth noting that the performance of polygenic models did not depend on the estimated degree of heritability, as evidenced by the absence of correlation between *GEV* and *H*^2^ (*ρ* = 0.04 for VSR, *ρ* = −0.05 for MLM, *p* > 0.05 in both cases). For instance, polygenic models allowed to explain around 45% of the broad-sense heritability, also for low-heritable traits, such as survival in Mediterranean sites, polycyclism, and SLA. *GEV* can be interpreted as an analogous of the SNP-based heritability, with the particularity that *GEV* refers to proportion of the variance in genetic values, rather than on the phenotypic values that are explained by associated SNPs (see Materials and Methods for further details). SNP-based heritability is becoming a fundamental parameter in quantitative genetics because it can yield insights into the ‘missing heritability’ of complex traits (Hou *et al.* 2019). In this sense, our study shows that polygenic approaches can be a promising strategy to account for a significant part of this missing heritability that is commonly observed in GWAS in forest trees (reviewed by Hall *et al.* 2016; Lind *et al.* 2018).

However, insights provided by SNP-based estimations of *GEV* should be interpreted with caution. First, because maritime pine has a huge genome size (around 28 Gbp; Grotkopp *et al.* 2004; Zonneveld 2012) and a rapid decay in linkage disequilibrium (Neale and Savolainen 2004), a larger number of genotyped SNPs should be needed to obtain a good genomic coverage. And second, because rare variants are usually difficult to incorporate in genotyping platforms, such as the one used in our study. Such rare variants may indeed account for an important proportion of the heritability in complex traits (Young 2019). Even though further investigations are needed to draw stronger conclusions, robust and consistent estimates of polygenicity across methods were fostered herein by a precise phenotypic evaluation in a large number of individuals (over 12,500 trees).

### Stability of polygenicity estimates across environments and years

The temporal and spatial heterogeneity of selection can impact the evolution of the genetic architecture underlying adaptation (Sella and Barton 2019). Monitoring the patterns of genetic architecture not only across environments but also across years is an important issue in long-lived forest trees that may experience changing selection pressures along their lifetimes. In this sense, our study is not only a validation of the polygenic adaptation model in a new organism, but a contribution to improving our understanding of adaptation. Surprisingly, the estimated degree of polygenicity remained stable across environments for all trait categories, especially tree height. Additionally, we observed highly stable genetic architectures for height, phenology, and survival across years. For the case of tree height, polygenicity was highly stable for three time-point measures along a time-span of 6 years, comprising seedling and juvenile stages, during which trees are more vulnerable and selection pressure are more pronounced (Leck *et al.* 2008). However, analysis of gene function enrichment (see below) suggests that different genetic pathways could be underlying phenotypic variation in contrasting environments. Moreover, differences in gene expression may also underlie adaptation under different environments and years (Mähler *et al.* 2017; Hämälä *et al.* 2020).

### The role of negative selection in polygenic adaptation

All significant correlations between SNP effect-size and MAF were negative (for tree height, bud burst and SLA), suggesting a genetic architecture modeled, at least partially, by the action of negative selection, i.e. SNPs with large effects are rare because they mostly have deleterious effects and are thus selected against (O’Connor *et al.* 2019). The MLM method did not allow elucidating whether negative estimates of *S* were the consequence of an enrichment of trait-increasing or trait-decreasing alleles (Zeng *et al.* 2018), but it certainly suggests that these traits have been under some form of negative selection. The effect of purifying selection is widespread in model plant genomes (Wright and Andolfatto 2008), and it has been largely evidenced in trees (Krutovsky and Neale 2005; Palmé *et al.* 2009; Eckert *et al.* 2013; De La Torre *et al.* 2017; Grivet *et al.* 2017). Indeed, negative selection, and its variation across populations and through time, has been pointed out as a main cause for maintaining polygenicity (Zeng *et al.* 2018; O’Connor *et al.* 2019). Thus, negative selection may also explain, at least partially, the degree of polygenicity observed for fitness-related traits in maritime pine (but see below), as well as the absence of large effect-size SNPs in previous association studies for this species (Lepoittevin *et al.* 2012; Budde *et al.* 2014; Hurel *et al.* 2019).

Nevertheless, strikingly, the negative selection patterns observed across environments and years did not mimic the trend observed for polygenicity. That is, negative selection was consistently inferred for height, but its strength changed across environments and years for survival and phenology-related traits. This uncoupling between negative selection and polygenicity may result from the fact that our limited coverage of maritime pine genome did not account for (most) rare variants, which can considerably affect *S* estimates (Zeng *et al.* 2018). In addition, polygenic adaptation generally results in highly stochastic genetic responses driven by non-predictable changes in allele frequencies (Zhang *et al.* 2013).

Finally, we detected signals of gene enrichment for 10 pathways that had higher values of maximum SNP effect-size or higher posterior probability of being included in the polygenic models: height in the French Atlantic environment and survival in the Iberian Atlantic environment were enriched for genes coding for *transcription factors*, bud burst in 2015 for genes within the *monolignol biosynthesis* pathway, and *D. sapinea* susceptibility (considering both the induced necrosis and needle discoloration) for genes within the *ubiquitin system*, *signal transduction* and *flavonoid biosynthesis* pathways. Assuming that evolution of these pathways is driven by negative selection, these patterns could be interpreted as a consequence of the accumulation of (slightly) deleterious alleles, resulting in higher proportions of SNPs with non-zero effect-size on these phenotypic traits. This higher tolerance to retain deleterious mutations could be explained by a high genetic redundancy (Nowak *et al.* 1997; Krakauer and Nowak 1999). Otherwise, if we were to assume a higher impact of positive than negative selection, the observed patterns would imply an accumulation of beneficial mutations in these pathways, which is a hypothesis worth exploring using sequence-based neutrality tests in future studies.

Another five pathways were enriched in lower effect-sizes alleles: genes involved in *cytoskeleton* were linked with height in the Mediterranean environment, those in the *glycan metabolism* pathway were associated with bud burst in 2015, and those for *cell growth and death*, *DNA recombination and repair*, and *UV response* were associated with phenology growth index. These pathways perform general functions and could be constituted by functionally important genes. In this case, the observed patterns suggest higher genetic constraints on these functionally important genes, for which negative selection should be highly efficient (Wright and Andolfatto 2008). Interestingly, our results suggest that even for stable estimates of polygenicity, different gene pathways could underlie polygenic adaptation for height in contrasting environments. Finally, although our gene enrichment analysis revealed some pathways with stronger evidence for polygenic adaptation, we cannot discard the influence of other (non-studied) gene pathways, as pointed by the omnigenic theory (Boyle *et al.* 2017).

## Conclusions

The study of genetic adaptation is currently facing new challenges. The advancement of GWAS relies on the development of methods able to detect causal variants of small effect-size, or at low allele frequencies. Our study, adopting a polygenic adaptation model on well-characterized maritime pine clones planted in contrasted environments, contributed to a better understanding of the heritability of complex adaptive traits in long-lived organisms, and its underlying genetic architecture. Our results showed that most complex adaptive traits are polygenic, with several of them showing also signatures of negative selection. The degree of polygenicity was similar for traits spanning different functional categories, and this genetic architecture was considerably stable over time and across environments. Current models for predicting population trajectories in forest trees under climate change are based on identification of outlier SNPs with relatively large effects on phenotypes and/or strong correlation with climate variables (e.g. Jaramillo-Correa *et al.* 2015; Rellstab *et al.* 2016; Lu *et al.* 2019). Because polygenic adaptation can take place rapidly (see, for example, Jain and Stephan 2017), current prediction models are probably underestimating the capacity of natural forest tree populations to adapt to new environments. Thus, adopting a polygenic adaptation perspective could significantly improve prediction accuracy, and provide new scenarios to inform forest conservation and reforestation programs (Valladares *et al.* 2014; Fady *et al.* 2016). Also, a better understanding of the genetic architecture of economically valuable polygenic traits can improve genomic-assisted breeding, and allow building better genomic selection models (Grattapaglia *et al.* 2018).

## Acknowledgements

We thank A. Saldaña, F. del Caño, E. Ballesteros and D. Barba (INIA) and the ‘Unité Expérimentale Forêt Pierroton’ (UEFP, INRAE; doi: 10.15454/1.5483264699193726E12) for field assistance. Data used in this research are part of the Spanish Network of Genetic Trials (GENFORED, http://www.genfored.es). We thank all persons and institutions linked to the establishment and maintenance of field trials used in this study. Thanks are extended to Antoine Kremer, Martin Lascoux and Outi Savolainen for valuable insights and discussions on models of local adaptation in forest trees.

## Funding

This study was funded by the Spanish Ministry of Economy and Competitiveness through projects RTA2010-00120-C02-02 (CLONAPIN), CGL2011-30182-C02-01 (AdapCon) and AGL2012-40151-C03-02 (FENOPIN). The study was also supported by the ‘Initiative d’Excellence (IdEx) de l’Université de Bordeaux - Chaires d’installation 2015’ (EcoGenPin) and the European Union’s Horizon 2020 research and innovation programme under grant agreement No 773383 (B4EST).

## Author contributions

Mde-M collected field data, carried out the statistical analyses and drafted the manuscript. JM, RA and CP designed and established the common gardens, and helped with field data collection. IR-Q, DG, CP, GGV and SCG-M contributed to the SNP assay design and molecular laboratory work. AE, RA and SCG-M conceived and designed the study. IR-Q and AH collected field data and helped with the statistical analyses. JPJ-C identified gene pathways and defined gene-sets. MH, SCG-M, DG, JPJ-C and GGV contributed to the statistical analysis of genomic data. SCG-M coordinated the study. All authors contributed to manuscript discussion and review, and gave final approval for publication.

## Supplemental material

**Table S1. Phenotypic data summary and quantitative genetic analysis**. *V*_g_ stands for genetic variance (posterior mean of the variance explained by clone effect), *H*^2^ stands for broad-sense heritability and *Q*_ST_ for genetic differentiation among populations (posterior mode and 95% credible interval are presented).

**Table S2. MCMCglmm Bayesian model parametrization**. Psrf stands for the Gelman-Rubin potential scale reduction factor criterion, a measure of model convergence. Good convergence of models is expected for psrf < 1.02.

**Table S3. List of genes included in the 17 gene sets considered for gene function enrichment at pathway level.** Annotation based on KEGG: Kyoto Encyclopedia of Genes and Genomes (https://www.genome.jp/kegg/) is also provided. *Annotation* label indicates genes for which no hit with KEGG database was found and thus were assigned to metabolic pathways/modules based on the InterPro annotation.

**Table S4. Number of non-zero effect-size SNPs (***nbnon-zero***) and genetic explained variance (***GEV***) estimated using Bayesian variable selection regression (VSR), as implemented in piMASS software, and the Bayesian linear mixed model, MLM, implemented in GCTB software.** For MLM, the coefficients of correlation between SNP effect-size and minor allele frequency (S) are also provided. The parameters are presented as the posterior median and 95% credible intervals. Estimates not overlapping zero are marked in bold. NA: models that did not converge.

**Figure S1. Sampled maritime pine populations (circles) and common garden sites (other symbols).** Neutral gene pools (identified in Jaramillo-Correa *et al.* 2015) outline the species natural distribution range in different colors.

**Figure S2. Posterior distribution of the number of non-zero size-effect SNPs for 26 traits belonging to five categories: survival, height, phenology-related, functional, and biotic-stress response traits.** The number of non-zero size-effect SNPs was estimated through two Bayesian methods: posterior inference via model averaging and subset selection (VSR), as implemented in the software piMASS (Guan and Stephens 2011), and the Mixed Linear Model (MLM) implemented in the software CGTB (Zeng *et al.* 2018). The posterior median is indicated with a dashed line.

**Figure S3. Posterior distribution of SNP effect-sizes for 26 traits belonging to five categories: survival, height, phenology-related, functional, and biotic-stress response traits.** SNP effect-size was estimated through two Bayesian methods: posterior inference via model averaging and subset selection (VSR), as implemented in the software piMASS (Guan and Stephens 2011), and the Mixed Linear Model (MLM) implemented in the software CGTB (Zeng *et al.* 2018).

## Notes

### Competing Interest Statement

The authors have declared no competing interest.

### Summary of Updates

Additional analysis on gene function enrichment at pathway-level have been incorporated. Writting of all sections has been reorganized and improved.

