## Supplemental Figure S1 for "Polygenic adaptation and negative selection across traits, years and environments in a long-lived plant species (*Pinus pinaster* Ait., Pinaceae)"

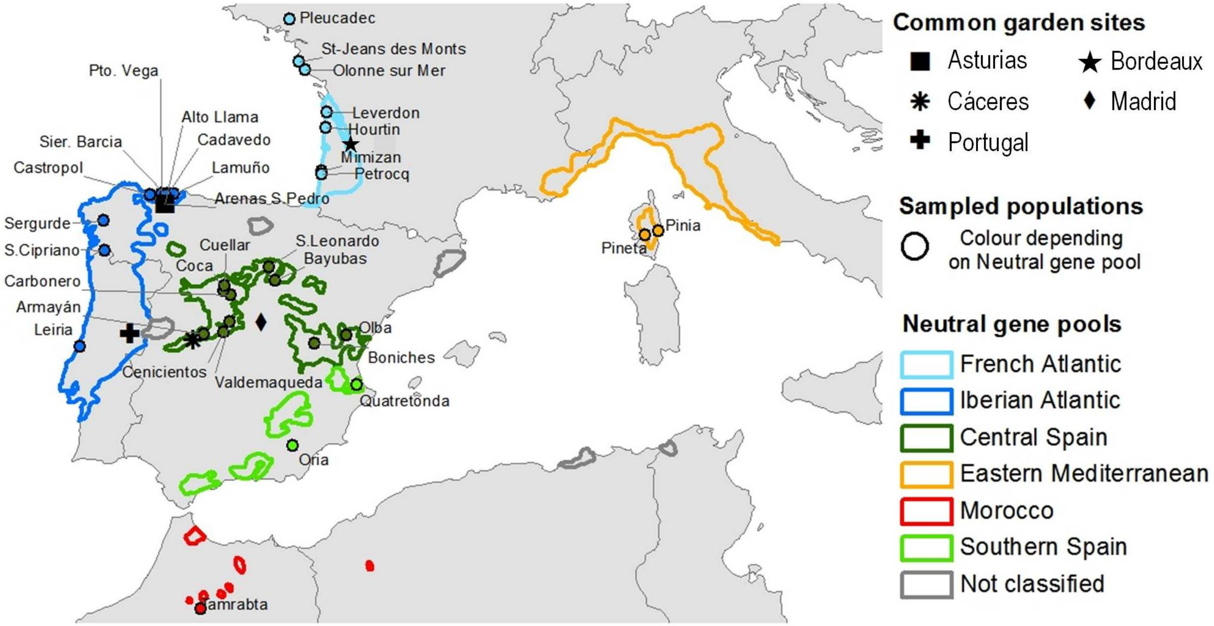


**Figure S1. Sampled *Pinus pinaster* populations (circles) and common garden trial sites (other symbols).** Neutral gene pools (identified in Jaramillo-Correa *et al.* 2015) outline the species natural distribution range in different colors .
