## Supplemental Figure S2 for "Polygenic adaptation and negative selection across traits, years and environments in a long-lived plant species (*Pinus pinaster* Ait., Pinaceae)"

**Figure S2. Posterior distribution of the number of non-zero size-effect SNPs for 26 traits belonging to five categories: survival, height, phenology-related, functional, and biotic-stress response traits.** The number of non-zero size-effect SNPs was estimated through two Bayesian methods: posterior inference via model averaging and subset selection (VSR), as implemented in the software piMASS (Guan and Stephens 2011), and the Mixed Linear Model (MLM) implemented in the software CGTB (Zeng *et al.* 2018). The posterior median is indicated with a dashed line.

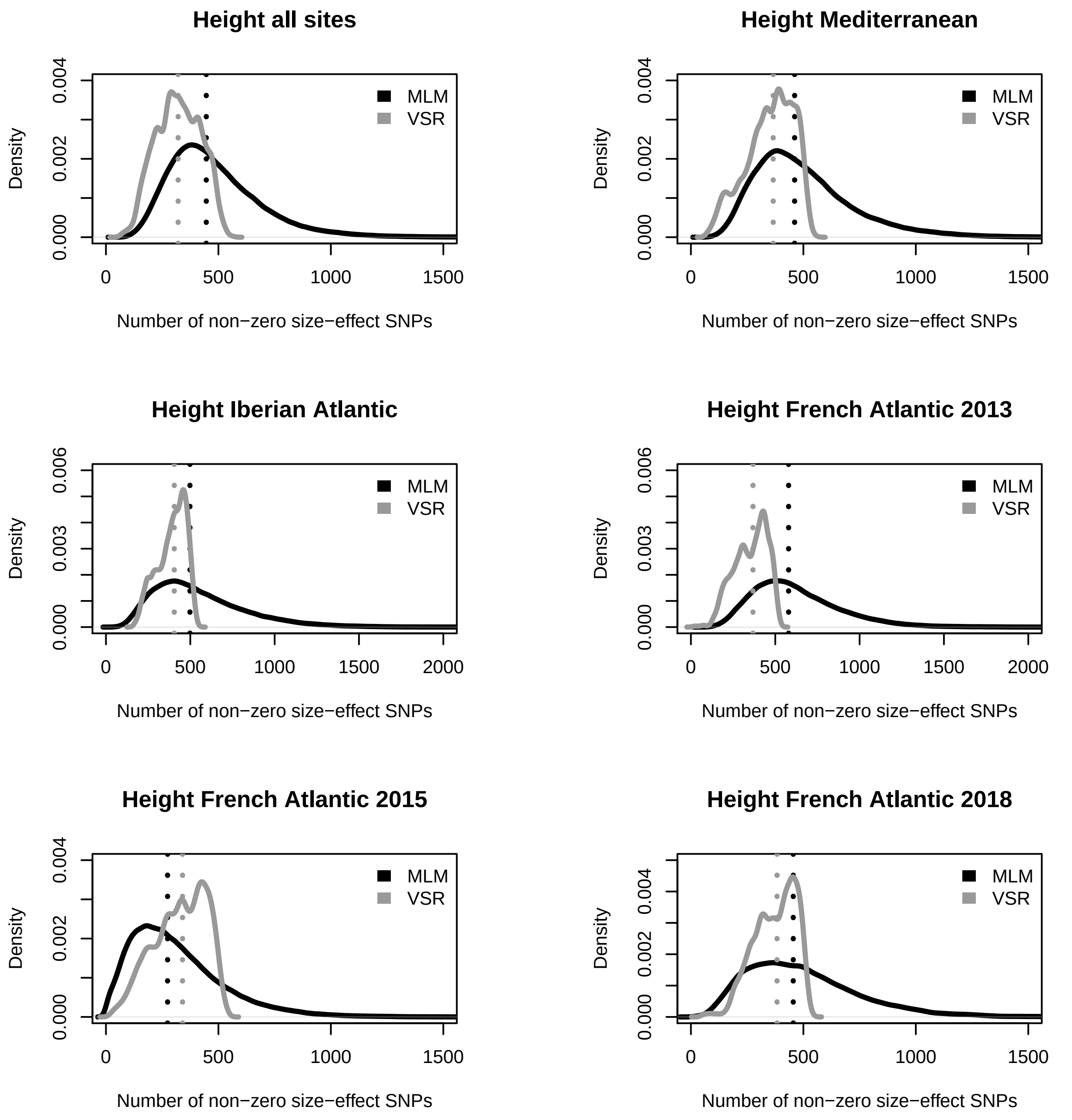
Height all sites
Height Mediterranean
Height Iberian Atlantic
Height French Atlantic 2013
Height French Atlantic 2015
Height French Atlantic 2018
Density
Number of non−zero size−effect SNPs
MLM
VSR
0.000
0.002
0.004
0.003
0.006
0
500
1000
1500
2000

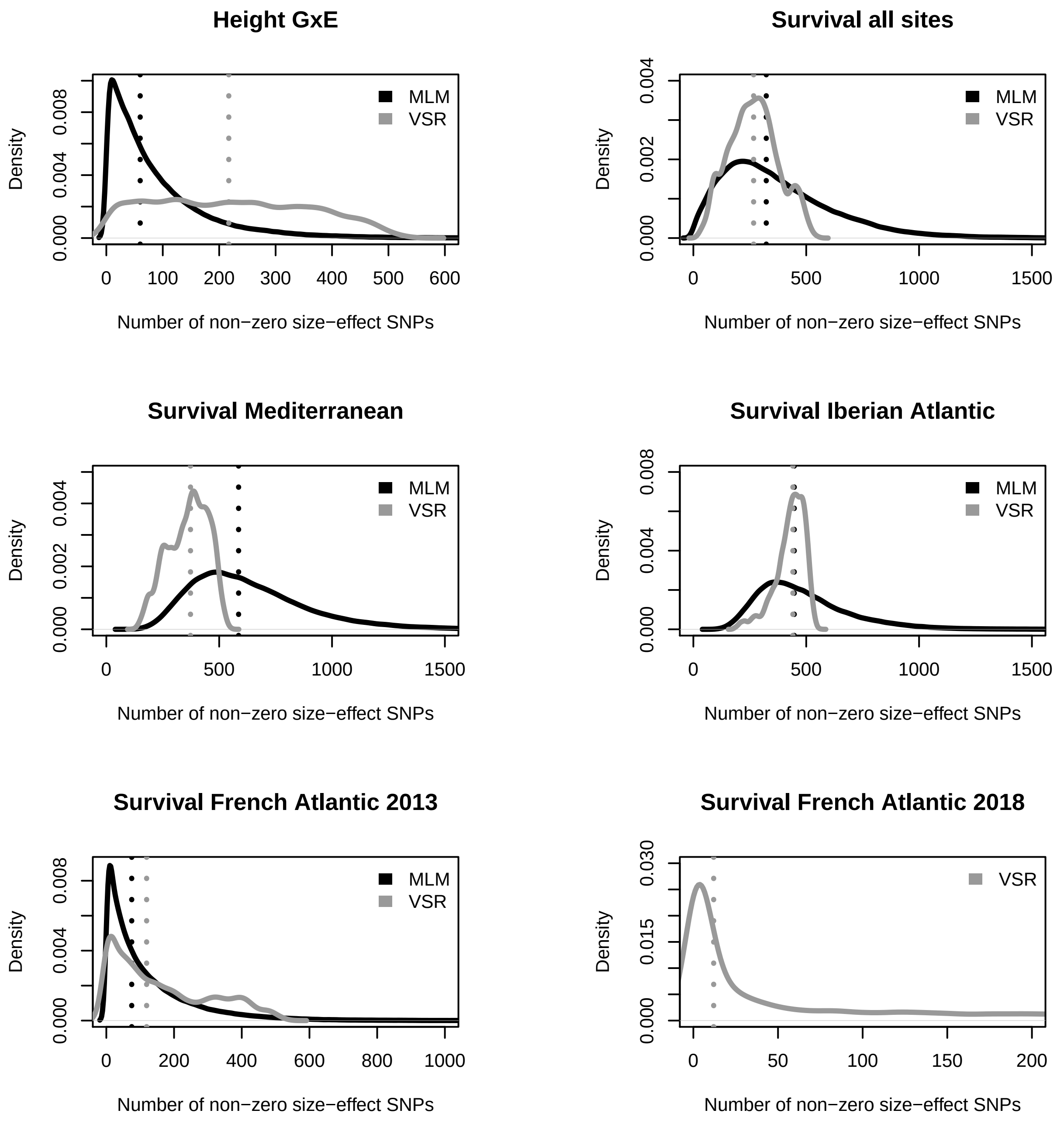
Height GxE
Density
0.000
0.004
0.008
0 100 200 300 400 500 600
Number of non−zero size−effect SNPs
MLM
VSR
Survival all sites
Density
0.000
0.002
0.004
0 500 1000 1500
Number of non−zero size−effect SNPs
MLM
VSR
Survival Mediterranean
Density
0.000
0.002
0.004
0 500 1000 1500
Number of non−zero size−effect SNPs
MLM
VSR
Survival Iberian Atlantic
Density
0.000
0.004
0.008
0 500 1000 1500
Number of non−zero size−effect SNPs
MLM
VSR
Survival French Atlantic 2013
Density
0.000
0.004
0.008
0 200 400 600 800 1000
Number of non−zero size−effect SNPs
MLM
VSR
Survival French Atlantic 2018
Density
0.000
0.015
0.030
0 50 100 150 200
Number of non−zero size−effect SNPs
VSR

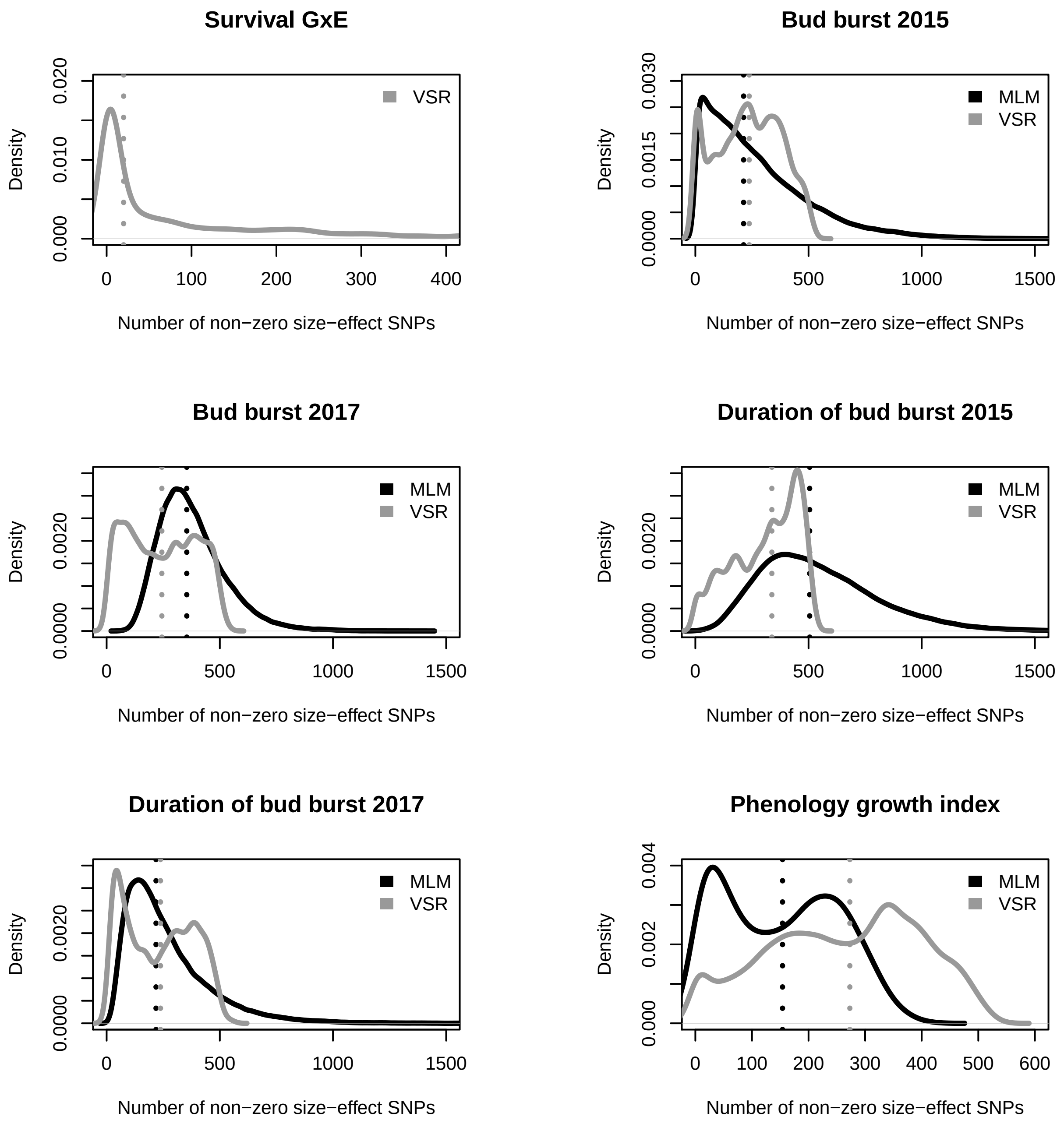
Survival GxE
VSR
Density
0.000
0.010
0.020
0
100
200
300
400
Number of non−zero size−effect SNPs
Bud burst 2015
MLM
VSR
Density
0.0000
0.0015
0.0030
0
500
1000
1500
Number of non−zero size−effect SNPs
Bud burst 2017
MLM
VSR
Density
0.0000
0.0020
0
500
1000
1500
Number of non−zero size−effect SNPs
Duration of bud burst 2015
MLM
VSR
Density
0.0000
0.0020
0
500
1000
1500
Number of non−zero size−effect SNPs
Duration of bud burst 2017
MLM
VSR
Density
0.0000
0.0020
0
500
1000
1500
Number of non−zero size−effect SNPs
Phenology growth index
MLM
VSR
Density
0.000
0.002
0.004
0
100
200
300
400
500
600
Number of non−zero size−effect SNPs

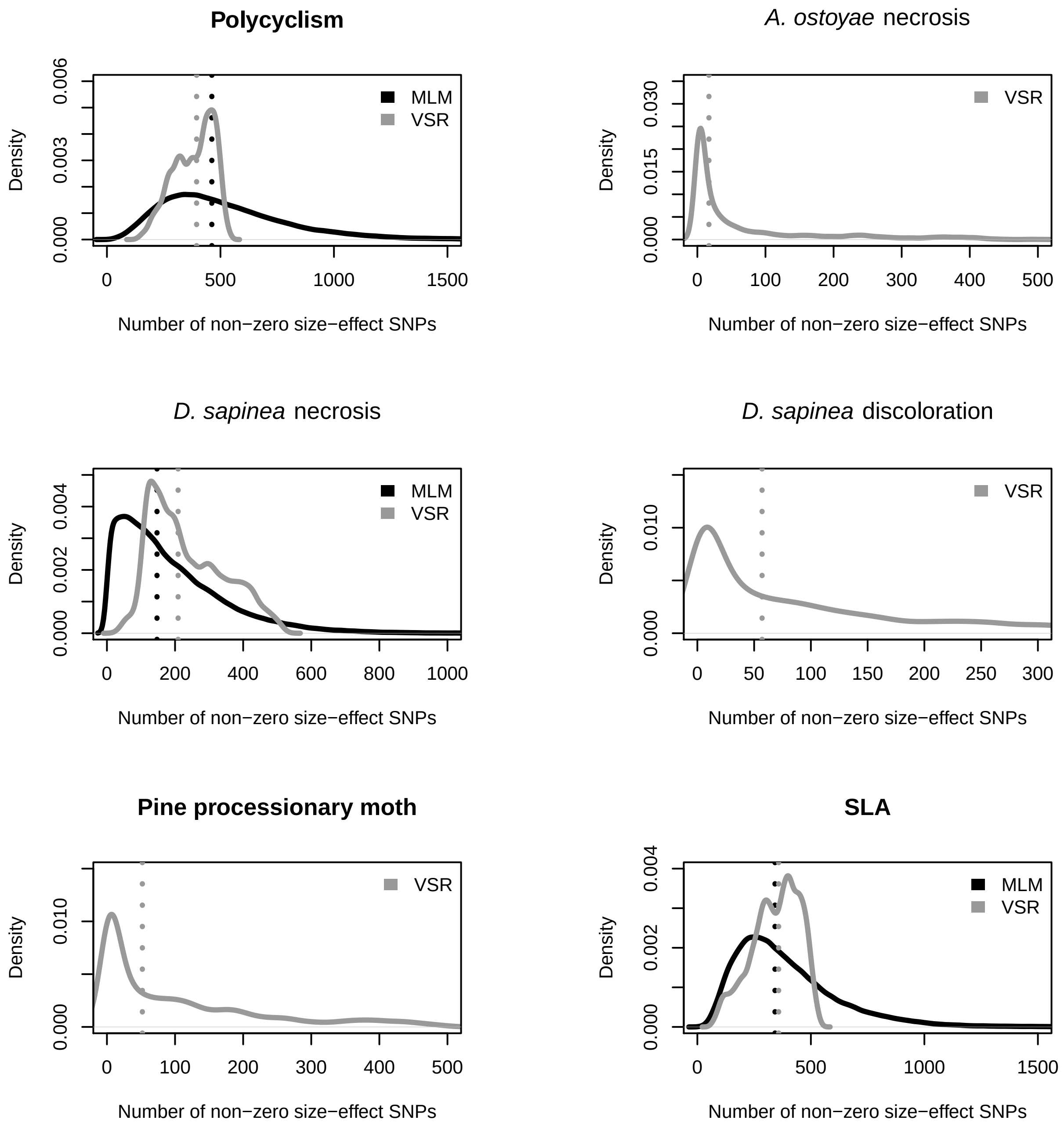
Polycyclism
MLM
VSR
Density
0.000
0.003
0.006
0
500
1000
1500
Number of non−zero size−effect SNPs
A. ostoyae necrosis
VSR
Density
0.000
0.015
0.030
0
100
200
300
400
500
Number of non−zero size−effect SNPs
D. sapinea necrosis
MLM
VSR
Density
0.000
0.002
0.004
0
200
400
600
800
1000
Number of non−zero size−effect SNPs
D. sapinea discoloration
VSR
Density
0.000
0.010
0
50
100
150
200
250
300
Number of non−zero size−effect SNPs
Pine processionary moth
VSR
Density
0.000
0.010
0
100
200
300
400
500
Number of non−zero size−effect SNPs
SLA
MLM
VSR
Density
0.000
0.002
0.004
0
500
1000
1500
Number of non−zero size−effect SNPs

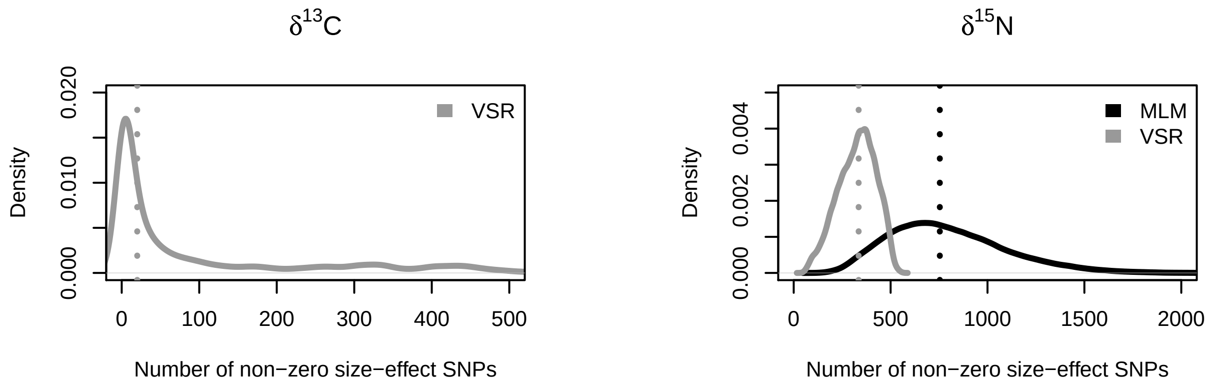
δ13C
Density
0.000
0.010
0.020
VSR
0
100
200
300
400
500
Number of non−zero size−effect SNPs
δ15N
Density
0.000
0.002
0.004
MLM
VSR
0
500
1000
1500
2000
Number of non−zero size−effect SNPs
