## Supplemental Figure S3 for "Polygenic adaptation and negative selection across traits, years and environments in a long-lived plant species (*Pinus pinaster* Ait., Pinaceae)"

**Figure S3. Posterior distribution of SNP effect-sizes for 26 traits belonging to five categories: survival, height, phenology-related, functional, and biotic-stress response traits.** SNP effect-size was estimated through two Bayesian methods: posterior inference via model averaging and subset selection (VSR), as implemented in the software piMASS (Guan and Stephens 2011), and the Mixed Linear Model (MLM) implemented in the software CGTB (Zeng *et al.* 2018).

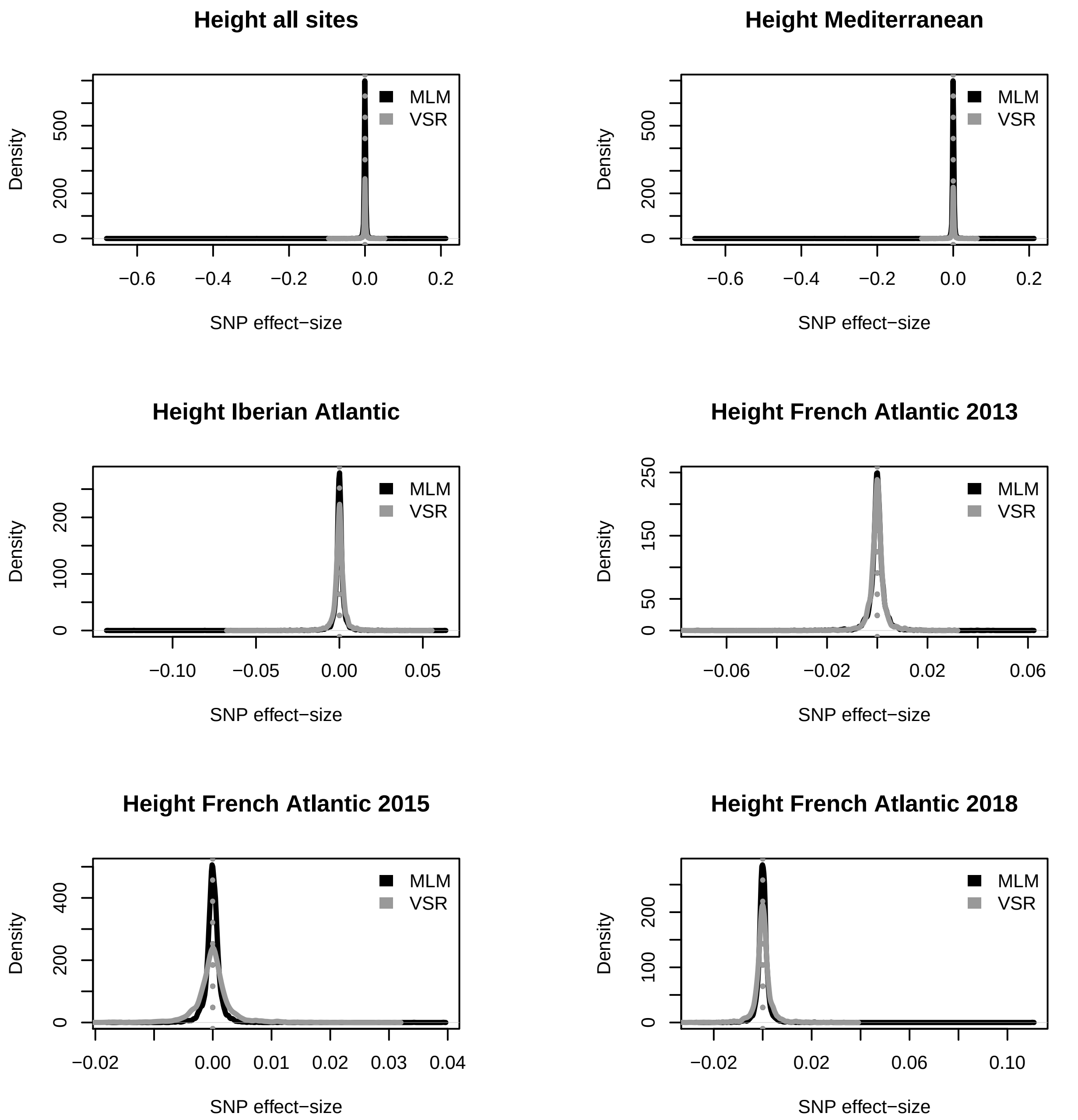
Height all sites
MLM
VSR
Density
0
200
500
−0.6
−0.4
−0.2
0.0
0.2
SNP effect−size
Height Mediterranean
MLM
VSR
Density
0
200
500
−0.6
−0.4
−0.2
0.0
0.2
SNP effect−size
Height Iberian Atlantic
MLM
VSR
Density
0
100
200
−0.10
−0.05
0.00
0.05
SNP effect−size
Height French Atlantic 2013
MLM
VSR
Density
0
50
150
250
−0.06
−0.02
0.02
0.06
SNP effect−size
Height French Atlantic 2015
MLM
VSR
Density
0
200
400
−0.02
0.00
0.01
0.02
0.03
0.04
SNP effect−size
Height French Atlantic 2018
MLM
VSR
Density
0
100
200
−0.02
0.02
0.06
0.10
SNP effect−size

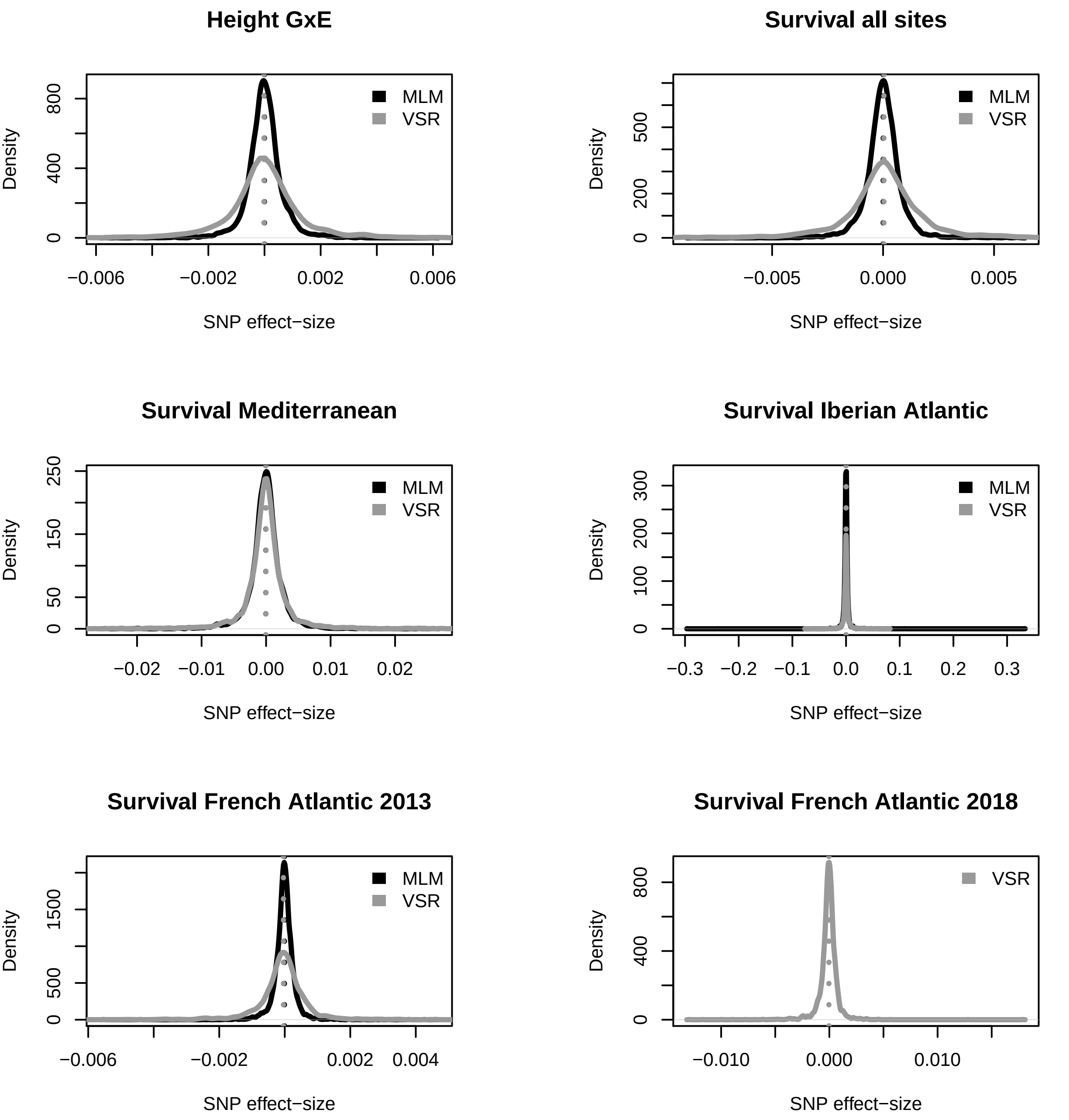
Height GxE
Density
0
400
800
MLM
VSR
−0.006
−0.002
0.002
0.006
SNP effect−size
Survival all sites
Density
0
200
500
MLM
VSR
−0.005
0.000
0.005
SNP effect−size
Survival Mediterranean
Density
0
50
150
250
MLM
VSR
−0.02
−0.01
0.00
0.01
0.02
SNP effect−size
Survival Iberian Atlantic
Density
0
100
200
300
MLM
VSR
−0.3
−0.2
−0.1
0.0
0.1
0.2
0.3
SNP effect−size
Survival French Atlantic 2013
Density
0
500
1500
MLM
VSR
−0.006
−0.002
0.002
0.004
SNP effect−size
Survival French Atlantic 2018
Density
0
400
800
VSR
−0.010
0.000
0.010
SNP effect−size

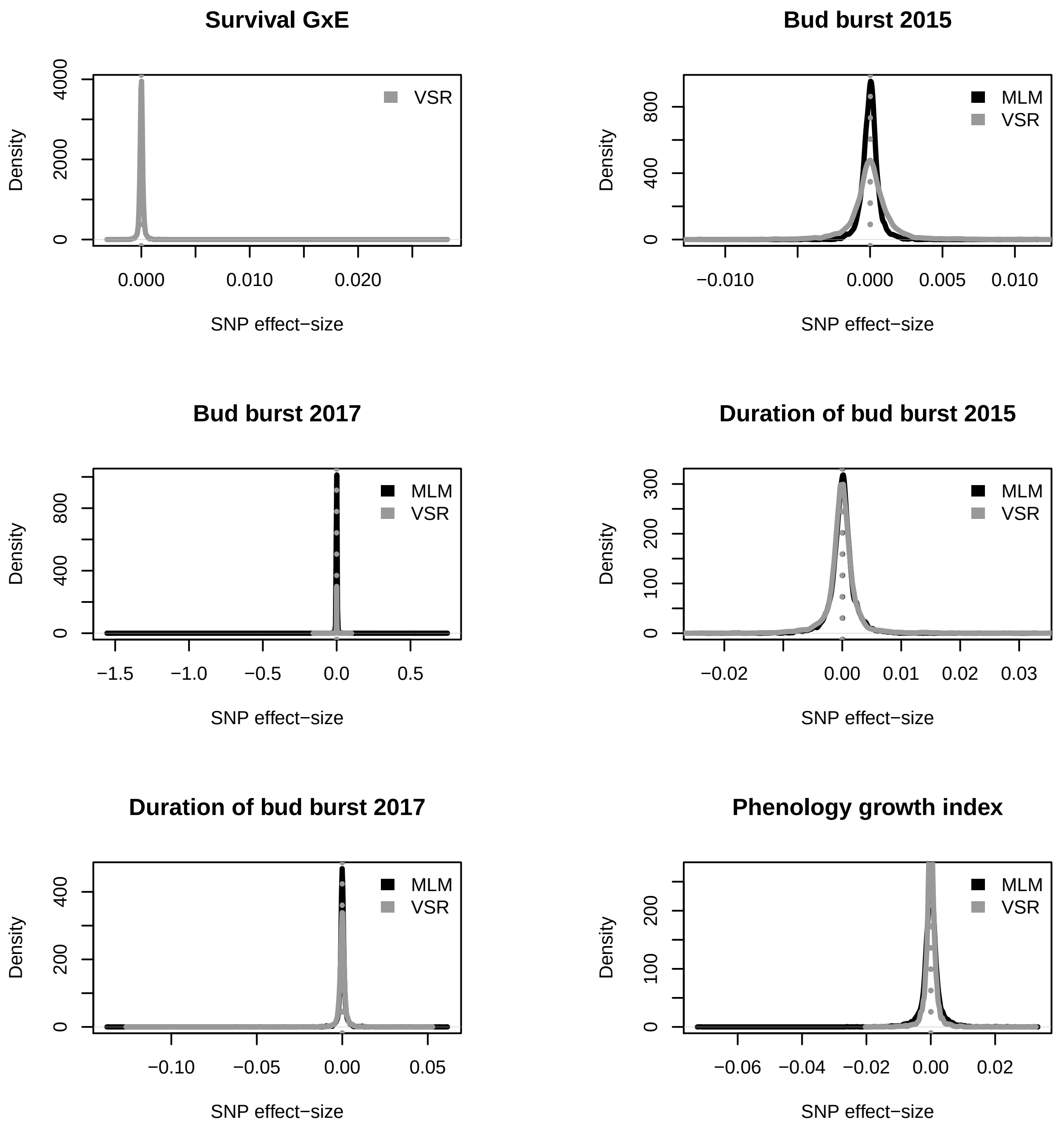
Survival GxE
VSR
Density
0
2000
4000
0.000
0.010
0.020
SNP effect−size
Bud burst 2015
MLM
VSR
Density
0
400
800
−0.010
0.000
0.005
0.010
SNP effect−size
Bud burst 2017
MLM
VSR
Density
0
400
800
−1.5
−1.0
−0.5
0.0
0.5
SNP effect−size
Duration of bud burst 2015
MLM
VSR
Density
0
100
200
300
−0.02
0.00
0.01
0.02
0.03
SNP effect−size
Duration of bud burst 2017
MLM
VSR
Density
0
200
400
−0.10
−0.05
0.00
0.05
SNP effect−size
Phenology growth index
MLM
VSR
Density
0
100
200
−0.06
−0.04
−0.02
0.00
0.02
SNP effect−size

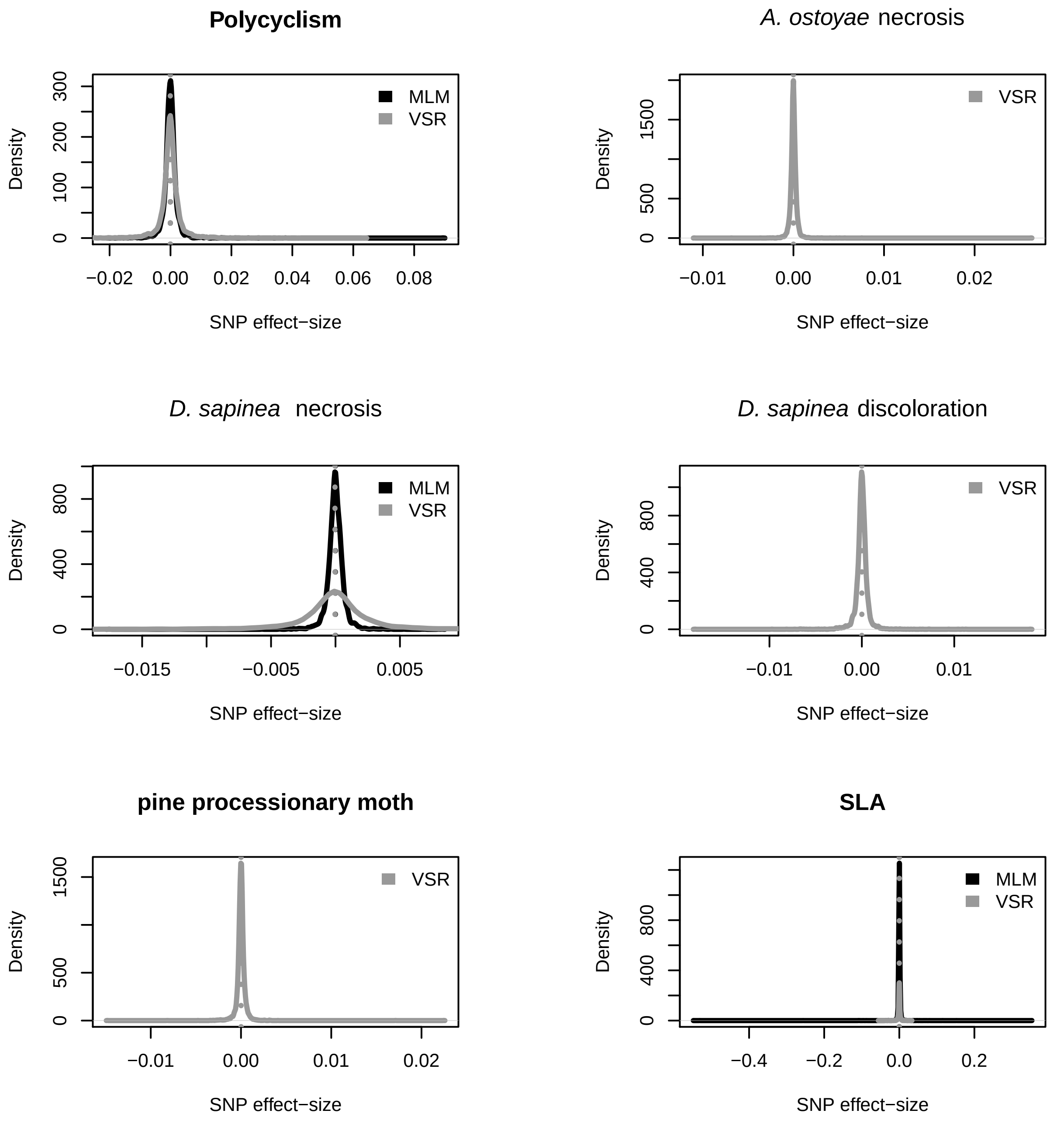
Polycyclism
Density
SNP effect−size
MLM
VSR
300
200
100
0
−0.02
0.00
0.02
0.04
0.06
0.08
A. ostoyae necrosis
VSR
1500
500
0
−0.01
0.00
0.01
0.02
D. sapinea necrosis
MLM
VSR
800
400
0
−0.015
−0.005
0.005
D. sapinea discoloration
VSR
800
400
0
−0.01
0.00
0.01
pine processionary moth
VSR
1500
500
0
−0.01
0.00
0.01
0.02
SLA
MLM
VSR
800
400
0
−0.4
−0.2
0.0
0.2

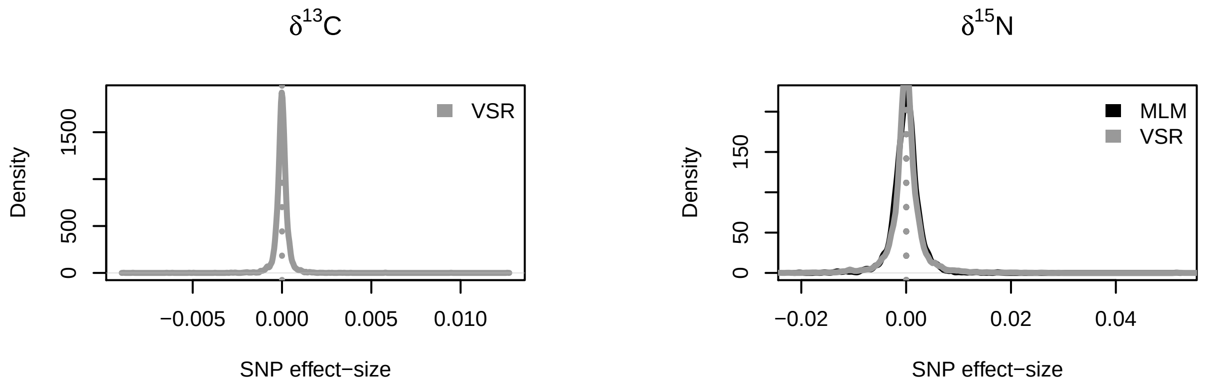
δ13C
VSR
1500
500
0
Density
−0.005
0.000
0.005
0.010
SNP effect−size
δ15N
MLM
VSR
150
50
0
Density
−0.02
0.00
0.02
0.04
SNP effect−size
